# PHF6 Interacts with LMO2 During Normal Haematopoiesis and in Leukaemia and Regulates Gene Expression and Genome Integrity

**DOI:** 10.1101/2020.08.18.255471

**Authors:** Vesna S. Stanulović, Sarah Binhassan, Ian Dorrington, Douglas G. Ward, Maarten Hoogenkamp

**Affiliations:** Institute of Cancer and Genomics Sciences, College of Medical and Dental Sciences, University of Birmingham, Birmingham B15 2TT, UK

## Abstract

The transcriptional mediator LIM domain only 2 (LMO2) forms a large multi-protein complex together with TAL1/LYL1, HEB/E2A, LDB1 and GATA. This complex regulates transcription from the onset of haematopoietic development and during differentiation. Chromosomal re-arrangements involving LMO and other members of the complex are causative for T-cell lymphoblastic leukaemia (T-ALL). We have identified Plant Homeodomain (PHD)-like Finger 6 (PHF6) as a new LMO2 interacting factor. Somatic mutations in *PHF6* have been found to occur in several types of leukaemia. We show that PHF6 interacts with LMO2 during the initial stages of the haematopoietic development, myeloid differentiation and in T-ALL. The LMO2/PHF6 complex binds the DNA and regulates linage-specific gene expression. Additionally, a loss or reduction of LMO2 and PHF6 leads to chromosomal instability. PHF6 and LMO2 are required for maintaining levels of γH2AX and 53BP1, where PHF6 is important for γH2AX accumulation and LMO2 has a role in recruiting 53BP1 to γH2AX foci.

## Introduction

LIM domain only 2 (LMO2) is a 17kDa protein, consisting of two LIM domains that are involved in protein-protein interactions. LMO2 functions in a transcriptional complex containing TAL1/LYL1, HEB/E2A, LDB1 and GATA factors to regulate many haematopoietic and non-haematopoietic genes [1–5]. The essential role of the LMO2 complex in haematopoiesis is facilitated through the ability of LDB1 to trimerise and facilitate long range interactions between distal regulatory DNA elements [6, 7]. LMO2 is expressed from the haemangioblast stage onwards, in haematopoietic stem and progenitor cells and downregulated during differentiation in all but the erythroid lineage [8–10]. Additionally, LMO2 was one of six proteins found to be required for reprogramming committed blood cells to induced haematopoietic stem cells (HSCs), further illustrating its importance for blood development [11].

During normal T-cell development, expression of both LMO2 and TAL1 is downregulated in CD4/CD8 double negative thymocytes. Recurrent chromosomal aberrations involving *LMO2* and *TAL1* cause these genes to remain expressed resulting in the development of T-cell Acute Lymphoblastic Leukaemia (T-ALL) [12–15]. Transgenic mice overexpressing LMO2 efficiently developed T-ALL, although with long latency [16], whereas additional overexpression of TAL1 reduced the latency [17]. Gene expression profiling analysis on paediatric T-ALL patient samples showed that cases with TAL/LMO rearrangements constitute a separate cluster which accounts for 45-50% of T-ALL [18, 19].

Human PHD finger protein 6 (PHF6) is a 365 amino acid protein that was first described as the gene whose inactivating germline mutations cause Börjeson-Forssman-Lehmann syndrome (BFLS; OMIM 301900) [20]. BFLS is a rare X-linked disorder, characterized by severe mental retardation, epilepsy, hypogonadism, obesity and facial dysmorphia, including narrow palpebral fissure and large ears. A role for somatic mutations of the PHF6 gene has been described in leukaemia. PHF6 mutations were found in 18% of paediatric and 36% of adult primary T-ALL patient samples, which were almost exclusively male [21]. In acute myeloid leukaemia (AML), 2-3% of cases harboured PHF6 mutations [22, 23], whereas PHF6 was mutated in 16-55% of mixed phenotype acute leukaemia (MPAL), based on relatively small cohorts [24–26]. Additionally, small numbers of PHF6 mutations were found in chronic myeloid leukaemia (CML) and high grade B cell lymphoma (HGBL) [27, 28]. Importantly, the mutations observed in BFLS are largely different to those found in leukaemia. In BFLS the mutations are either missense mutations that occur throughout the coding region or a nonsense mutation, 23 amino acids before the stop codon. The mutations identified in leukaemia are predominantly nonsense and frame shifts, with missense mutations accounting for less than one third of all PHF6 mutations [29]. As with BFLS, mutations in leukaemia are spread throughout the coding region.

PHF6 consists of two atypical plant homeodomains (aPHD1/2) with nuclear and nucleolar localization sequences and is 97% identical between mouse and human [20]. A number of interactions of PHF6 with other proteins have been described. PHF6 has been reported as a negative regulator of ribosomal RNA synthesis through a direct interaction with upstream binding factor (UBF) involving its aPHD1 domain [30]. The aPHD2 domain can bind DNA in vitro, albeit without sequence specificity, and is involved in recruitment of the nucleosome remodelling and deacetylation (NuRD) complex through direct interaction with retinoblastoma-binding protein 4 (RBBP4) [29, 31]. PHF6 was also shown to interact with the polymerase-associated factor 1 (PAF1) complex with a role in neuronal migration [32]. Additionally, a role of PHF6 was identified in non-homologous end joining (NHEJ) and G2 checkpoint recovery [33]. These roles and interactions of PHF6 were all shown in non-haematological cell types.

Recently, several studies have addressed the function of PHF6 in the haematopoietic system through conditional PHF6 knock out systems [34–37]. Haematopoietic *Phf6* knock out was conducted prior to HSC emergence through crosses with Vav1-CRE or Tie2-CRE mouse lines. The studies concluded that *Phf6* knock out cells exhibited higher reconstitution capability compared to those of wild type (wt) mice and a reduction in double negative (DN) T-cells. However, other findings were not consistent between these four publications. Three out of four studies observed an increase in LSK (Lin^-^;Sca^+^;Kit^+^) cells and a different set of three out of four observed no significant difference in LT-HSCs (LSK;CD150^+^;CD48^-^), whereas for both findings the fourth research group concluded the opposite. Additionally, one group reported a reduction in granulocyte-macrophage progenitors (GMPs), whereas another reported an increase in common lymphoid progenitors (CLPs); however, these observations were not in agreement with those of the other research groups. When leukaemia development was assessed, two of the groups reported that loss of PHF6 lowered the threshold for NOTCH1-induced T-ALL, whereas one reported no synergism. The lack of consistency between the different studies extended to their transcriptomic analyses of *Phf6* knock out cells. Different findings included enrichment of TNFα signalling associated genes, interferon α/β signalling signature, cell cycle and differentiation. The transcriptomic variability was highlighted in one of the studies [35], which used three different CRE mouse strains for deleting *Phf6*. Upon RNAseq, they reported expression profiles that were specific to the CRE deleter strain or developmental stage, with very little overlap between the data sets.

PHF6 mutations were found to be positively associated with overexpression of the two homeobox transcription factors TLX1 and TLX3, but negatively associated with the TAL1/LMO2 subgroup of T-ALL [21, 38]. The latter observation is of particular interest as this could be the consequence of a functional relationship between the LMO2 complex and PHF6. Indeed, in our study we show the physical interaction between LMO2 and PHF6 in T-ALL, ES cell-derived haemangioblasts and myeloid progenitors.

## Materials and methods

### Cell culture

Human T-ALL cell lines (ARR, DU.528, HSB2, and CCRF-CEM) were cultured in RPMI 1640 medium (Gibco), supplemented with 10% heat inactivated Foetal bovine serum (HI-FBS, Life Technologies), 2 mM GlutaMax (Gibco), 100 units/ml penicillin, 100 μg/ml streptomycin and maintained at a concentration between 0.4 - 2.0 x 10^6^/ml.

The murine PUER cell line and its derivatives were maintained in phenol red-free Iscove’s Modified Dulbecco’s Medium (Gibco), supplemented with 10% HI-FBS, 2 mM GlutaMax, 100 units/ml penicillin, 100 μg/ml streptomycin and 10 μg/ml IL-3 (PeproTech, 213-13). For the initiation of terminal differentiation, medium was supplemented with 100 nM 4-hydroxy-tamoxifen (OHT) and cells were harvested at the indicated time points.

Mouse *Phf6* knock out ES cells were purchased from The KOMP repository (Phf6tm1b). Wild type CCB and *Lmo2* knock out cells and ES cell culture conditions have been described before [4]. Briefly, ES cells were maintained on a layer of primary mouse embryonic fibroblasts (MEFs) in KnockOut DMEM (Life Technologies), supplemented with 15% EScult FBS (Stemcell technologies), 100 U/ml penicillin, 100 μg/ml streptomycin, 25 mM Hepes buffer, 2.5 mM Glutamax (Life Technologies), non-essential amino acids (Sigma), 0.15 mM monothioglycerol (MTG), 10^3^ U/ml ESGRO (Millipore). Differentiation was initiated by diluting single cell suspensions to 3.5×10^4^ cells/ml in *in vitro* differentiation (IVD) medium, consisting of IMDM, supplemented with 15% HI-FBS, 100 U/ml penicillin, 100 μg/ml streptomycin, 0.15 mM MTG, 50 μg/ml ascorbic acid, 180 μg/ml human transferrin (Roche). Flk-1^+^ cells were isolated by magnetic cell sorting (MACS) using biotinylated Flk-1 antibody (eBioscience 13-5821), anti-biotin microbeads (Miltenyi Biotec 130-090-485) and MACS LS columns (Miltenyi Biotec 130-042-401) according to manufacturer’s instructions. For further differentiation of Flk1+ cells into haemogenic endothelium and haematopoietic progenitors, cells were cultured in gelatine-coated 8-well Lab-Te II Chamber Slide System (Thermofisher) in IMDM, supplemented with 10% HI FBS (Gibco 10500), 20% D4T conditioned medium (24), 100 units/ml penicillin, 100 g/ml streptomycin, 0.45 mM MTG, 25 g/ml ascorbic acid, 180 g/ml human transferrin (Roche), 5 ng/ml VEGF (PeproTech 450-32), 10 ng/ml IL-6 (PeproTech 216-16). Cell populations were analysed using Flk1-APC, c-Kit-APC, Tie2-PE and CD34-FITC antibodies (eBioscience 17-5821-81, 17-1171, 12-5987, 25-0411, 11-0341-82) by flow cytometry (Beckman Coulter CyAn ADP) or confocal microscopy using Confocal Microscope Zeiss LSM 880 with Airyscan.

PlatE cells were grown in DMEM (Gibco), supplemented with 10% HI-FBS, 2 mM GlutaMax, 100 U/ml penicillin, 100 μg/ml streptomycin. Cells were passed when reaching 70-80% confluence.

All cell lines were tested for mycoplasma contamination and incubated at 37° C in a humidified incubator with 5% CO2.

### Creation shPHF6 PUER cell line

*Phf6* shRNA sequences were designed using http://cancan.cshl.edu/RNAi_central/RNAi.cgi?type=shRNA and purchased from Sigma-Aldrich [39]. Designed shRNA (shPhf6_1 AACTGTGCATTGCATGATAAAG, shPHF6_2 AACAAGGAATGTGGACAGTTAC, shPhf6_3 CCCACATCCTCCCATGGAACAG, shPhf6_4 CACTCGGAAGCTGATTTAGAAG, shPhf6_5 CGGACAGTTACTGATATCTGAA, shPhf6_6 CAGAGGGAAATTGCATATATTT, shLmo2_1 AGCCGCCTGTCAGAAGCATTTC, shLmo2_2 CCCAGCCCTTAGAGAGAATTTA, shLmo2_3 ACCATAGTAACTGACAAGATTA) were embedded into mir30, cloned into pMSCVhygro and tested as described previously[40]. PlatE packaging cells were used to produce virus. As an initial test of the different shRNA sequences, PlatE cells were co-transfected with MigR1 expression plasmid containing *Phf6* or *Lmo2* cDNA followed by an IRES GFP sequence, and pMSCVhyg shRNA constructs, targeting firefly luciferase transcripts (shCtrl; negative control), GFP (shGFP; positive control), or the different shPhf6 or shLmo2 targeting constructs.

shPhf6 and shLmo2 PUER cells were generated by transfection of PlatE cells with the shPhf6 and shLmo2 pMSCVhyg constructs. Virus containing media was collected and PUER cells were transduced by spin-infection in the presence of 12 μg/ml polybrene (Merck) at 850 g for 2h. Cells with integrated shRNA were selected with 300μg/ml Hygromycin B (Invitrogen). The GFP expressing cells were isolated by FACS sorting (BD FACSAria Fusion).

### Protein extraction

Crude nuclear extracts were prepared by incubating 2×10^7^ cell/ml hypotonic buffer (10 mM HEPES pH7.6, 10 mM KCl, 1.5 mM MgCl_2_) for 30 min incubation on ice, followed by centrifugation. Nuclei were then resuspended at 3×10^8^ nuclei/ml hypertonic buffer (20 mM HEPES pH7.6, 420 mM NaCl, 1.5 mM MgCl2, 0.2 mM EDTA, 0.5% NP40, 20% Glycerol) and incubated 20 min on ice. Supernatant was collected by centrifugation and diluted with 1.8 volume of no-salt buffer (20 mM HEPES pH7.6, 1.5 mM MgCl2, 0.2 mM EDTA, 0.5% NP40, 20% glycerol). All buffers were supplemented with 1:1000 Phosphatase Protease Inhibitor (PPI, Roche). Protein concentration of the nuclear extracts was determined using a BCA Protein Assay Kit (Pierce Biotechnology).

### Co-Immuno Precipitation

Protein G coated Dynabeads (30 μg/μl, Life Technologies) were resuspended in PBS containing 3% BSA (Sigma) and 1 μg IgG or specific antibody per 10 μl beads, followed by incubation for 30 min on a rotator at 4° C. Antibody-coated beads were incubated with nuclear extract (100 μg NE per 10 μl beads) for 2h at 4° C. Following two washes samples were analysed by Western blotting or mass spectrometry. Input controls consisted of 100 μg of nuclear extract.

### Western blotting

Proteins were separated on 4–12% polyacrylamide gels (Life Technologies), transferred to nitrocellulose membranes, blocked and incubated overnight with primary antibody. Following three washes with PBS, the membranes were incubated for 2 h with the appropriate secondary antibody. Primary antibodies (listed in cellular stainings section) were used at a final concentration of 1 μg/ml and secondary antibodies were IRDye 680RD or 800RD (Li-Cor), which were used at a 1:2000 dilution. Fluorescence was detected with Odyssey CLx Imager (Li-Cor) using Odyssey v3.0 software.

### Mass spectrometry

Immunoprecipitation was performed as described above, using 100 μl beads, 10 μg IgG or specific antibody and 1 mg nuclear extract. Proteins were separated on 4-12% polyacrylamide gels and stained with Coomassie blue. Lanes were excised and divided into 12 slices. Gel slices were destained in 50% acetonitrile, 50 mM ammonium bicarbonate, equilibrated in 10% acetonitrile, 50 mM ammonium bicarbonate, reduced with 50 mM DTT (30 min at 56° C) and alkylated with 100 mM iodoacetamide (30 min at RT in the dark). After washing in 10% acetonitrile, 50 mM ammonium bicarbonate the gel-pieces were lyophilised and in-gel digestion performed by incubation with sequencing-grade trypsin (Promega) in 10% acetonitrile, 50 mM ammonium bicarbonate overnight at 37° C. Peptides were extracted with 1% formic acid, 10% acetonitrile and lyophilised. Peptides were dissolved in 1% formic acid and analysed by LC-MS/MS using an Impact ESI-Q-TOF-MS (Bruker Daltonics). Data were searched against Swiss-Prot human and mouse sequence databases using MASCOT and requiring peptide scores >25 and protein FDR <1%.

### Expression analysis

RNA was isolated using a NucleoSpin RNA kit (Macherey-Nagel) according to the manufacturer’s protocol, after which the concentration and quality were determined with a NanoDrop 2000 UV-Vis Spectrophotometer (Thermo Scientific). For cDNA synthesis, typically 2 μg RNA was reverse transcribed using Oligo(dT)12–18 primer and SuperScript II Reverse Transcriptase (Life Technologies). Gene expression was measured by quantitative PCR (qPCR), using Luna qPCR Master Mix (NEB M3003) on an ABI 7500 Real-Time PCR. Primers were: PHF6_5’ CAAAGGAAGGCCAAGAAAAA, PHF6_3’ CCCTACATGGCAAAATCCAC, GAPDH_5’ CCTGGCCAAGGTCATCCAT, GAPDHR_3’ AGGGGCCATCCACAGTCTT. Quantitation was carried out using a standard curve generated by serial dilution of cDNA. RNAseq libraries were prepared using the TruSeq Stranded mRNA Sample Preparation Kit (Illumina), using 4 μg total RNA per sample. Libraries were run on an Illumina NextSeq 500 sequencer obtaining 150nt paired-end (300 cycles) reads.

### Chromatin Immunoprecipitation (ChIP)

ChIP assays were performed as described [4]. Cells were crosslinked with 1% formaldehyde for twelve minutes at room temperature. Nuclei were isolated and the chromatin was sonicated for 15 cycles of 30s on 30s off at 4° C, using a Bioruptor (Diagenode). Immunoprecipitation was carried out using antibodies bound to protein G magnetic beads (30 μg/μl, LifeTechnologies). After elution, crosslinks were reversed overnight at 65° C, after which the DNA was isolated using Ampure PCR purification beads (Beckman). The following antibodies were used: goat anti-TAL1 (sc-12984X), mouse anti-PHF6 (sc-365237) (Santa Cruz), rabbit anti-Ldb1 (ab96799, Abcam), goat anti-LMO2 (AF2726), goat anti-GATA2 (AF2046) (R&D systems). The obtained material was analysed by qPCR on an ABI 7500 real-time PCR System using primers Chr18_5’ ACTCCCCTTTCATGCTTCTGATATCCATT, Chr18_3’ AGGTCCCAGGACATATCCATT, Runx1Enh_5’ AACTGCCGGTTTATTTTTCG, Runx1Enh_3’ TCTCTGGGAAGCCTCTTGAC for T-ALL cell lines; Chr2_5’ AGGGATGCCCATGCAGTCT, Chr2_3’ CCTGTCATCAGTCCATTCTCC, PU1 Enh_5’ GCTGTTGGCGTTTTGCAAT, PU1 Enh_3’ GGCCGGTGCCTGAGAAA for Flk-1^+^ and PUER cells. RNAseq libraries were prepared using the Illumina sample preparation protocol with indexed primers. Libraries were run on an Illumina NextSeq 500 obtaining 75nt single-end reads.

### Cellular Staining

ES cell-derived haemogenic endothelium cultured on gelatine-coated 8-well Lab-Tek II Chamber Slide System (ThermoFisher) was fixed using 4% PFA for 10 min, washed with PBS and blocked with 1% FBS for 30 min at RT, followed by fluorescent-immunostaining using c-Kit-APC, Tie2-PE and CD34-FITC antibodies (eBioscience 17-1171, 12-5987, 11-0341-82).

T-ALL cell lines, PUER cells and its derivatives were harvested and fixed in 4% PFA for 10 min at RT, washed with PBS, permeabilised with 0.05% Triton X-100 in PBS for 10 min and blocked with 1% FBS for 30 min at RT. Fixed cells were incubated with the indicated primary antibody 1 μg/ml for 1hr, followed by washes with PBS and incubation with the secondary antibody for 30 min. After washing with PBS cells were spun onto microscopy slides using a Cytospin III centrifuge (Shandon) at 500rpm for 2min. Nuclei were counterstained with DAPI. Primary antibodies used were goat-anti TAL1 (Santa Cruz, sc-12984X), rabbit-anti PHF6 (Bethyl Laboratories, A301-451A), goat-anti LMO2 (R&D systems, AF2726), rabbit-anti 53BP1 (Novus Biologicals, NB-100 904) and mouse-anti γH2A.X (pS139) (Abcam, ab26350). Secondary antibodies were Alexa Fluor anti-goat 488, anti-rabbit 488 or 568 and anti-mouse 647 (A11055, A21206, A10042, A21463, Molecular Probes). Confocal images were obtained using the same settings for the laser intensity and gain per experiment on a Carl Zeiss Confocal Microscope LSM 880 with Airyscan (Carl Zeiss, Germany). Dynamic range of the images was adjusted using Zeiss Zen (blue edition) software and the same parameters were used for all images within an experiment.

After cytospin, PUER cells were air-dried, fixed for 30 s, stained in using Kwik-Diff (9990700-Thermo Fisher). Slides were submerged into fixative, eosinophilic and basophilic stain for 30 s, washed in water and covered. Images were obtained on Leica DM6000 microscope using LAXS software.

### Proximity ligation assay (PLA)

Cells were harvested and fixed in 4% PFA for 10 min at RT, washed with PBS, permeabilised with 0.05% Triton X-100 in PBS for 10 min and blocked with 1% FBS for 30min at RT. PLA was conducted using a Duolink PLA kit (Sigma-Aldrich) according to the manufacturer’s protocol. Briefly, cells were incubated with PLA secondary probes conjugated with oligonucleotides (anti-rabbit PLUS and anti-goat MINUS) for 1 h at 37° C, followed by incubation with ligase for 30 min at 37° C. Cells were then transferred to microscopy slides using a Cytospin III centrifuge (Shandon). DNA amplification by rolling-circle was performed for 2 h at 37° C. After final washing steps, the cells were mounted with DAPI and analysed the same as immune-fluorescent staining.

### Cytogenetic analysis

Mitotic cells were arrested at metaphase by treatment with 20 ng/ml Colcemid (KryoMax, Gibco) for 2 h at 37° C. They were then pelleted at 300 g, 8 min and swelled by hypotonic treatment with 75 mM KCl for 20 min at 37° C. After this, cold fixative (3:1 methanol/glacial acetic acid) was added and they were pelleted at 300 g, 8 min. Cells were further fixed for 15 min at RT and then dropped onto humidified, chilled glass slides. The mitotic index, quality of metaphase spread, presence of cytoplasm and overlaps were evaluated under a phase contrast microscope. The slides were then dried in a humidified-bed and stained with Giemsa–modified solution (Gibco) for 3 min and rinsed with distilled water for 20-30 s. Cells were photographed and chromosomes were counted using a light microscope (Leica DM6000).

### Bioinformatics analysis

Global analysis of the genome wide data was performed on usegalaxy.org and usegalaxy.eu [41] and on the University of Birmingham High Performance Computing cluster. Acquired RNAseq reads were mapped to the mouse genome (GRCm38/mm10) using HISAT2 [42]. Transcripts were assembled using Cufflinks 2.0.0 based on the reference genome with quartile normalisation and effective length correction. A combined gtf file was produced using Cuffmerge 2.1.1 and used for determination of differential gene expression using Cuffdiff 2.0.1. Heat maps of differentially expressed genes and the hierarchical clustering, based on Pearson correlation with average linkage clustering, were computed by MultiExperiment Viewer v4.9.0. Gene ontology enrichment was performed using DAVID 6.8 and GREAT [43, 44]. GO terms for biological processes with *p*-value <0.05 were considered significant; categories with redundant terms were filtered out.

For ChIPseq analysis, reads were mapped to the mouse genome mm10 (GRCm38) using HISAT2. Peaks were called using MACS 2 and the resulting BED files were used for calculating the number of overlapping peaks [45, 46]. The matrix files for overlay were made by sorting the peaks according to the peak intensity score [47]. For ChIPseq integration with RNAseq, we used gene expression files that contained genes with FPKM values > 1 in at least one of the samples and the nearest 5’ and 3’ genes were selected based on the genomic coordinates of the ChIPseq peaks. Overlays were generated using EaSeq [48]. Genome wide data reported in this study are available from the NCBI Gene Expression Omnibus portal (GEO: GSE154675).

## Results

### PHF6 is an LMO2 binding partner at the onset of haematopoietic differentiation and in T-ALL

Our aim was to identify novel LMO2 interacting partners and to elucidate their function. Immunoprecipitation using αLMO2 antibody and nuclear protein extracts from Flk-1^+^ haemangioblasts and four different T-ALL cell lines (ARR, DU.528, HSB2 and CCRF-CEM,) followed my mass spectrometry identified PHF6 in four out of five samples (Supplementary file 1). We decided to explore the PHF6-LMO2 interaction, as mutations in PHF6 commonly occur in T-ALL and AML patient samples [29]. They are significantly negatively correlated with TAL/LMO2 chromosomal rearrangements [21, 38], implying that intact PHF6 function may be required for TAL/LMO2-induced leukaemogenesis.

### PHF6 interacts with LMO2 in T-ALL cell lines

RNAseq data and qPCR showed that *PHF6* mRNA is expressed in all four T-ALL cell lines while western blot confirmed the presence of PHF6 protein (Figure 1A,B,C). Furthermore, immunofluorescent staining showed overlapping nuclear localisation of PHF6 and LMO2 (Figure 1D). Co-immunoprecipitation of LMO2 and PHF6 followed by western blotting confirmed the PHF6-LMO2 interaction (Figure 1E). Additional validation of the interaction was achieved in proximal ligation assays combining αLMO2 and αPHF6 antibodies in HSB2 and CCRF-CEM cells (Figure 1F). The combination of αLMO2 and αLDB1 or IgG was used as technical positive and negative control, respectively, and as a biological negative control we performed the same assay on Jurkat T-ALL cells, which express LMO1 instead of LMO2. Taken together, we conclude that PHF6 interacts with LMO2.

**Figure 1.**
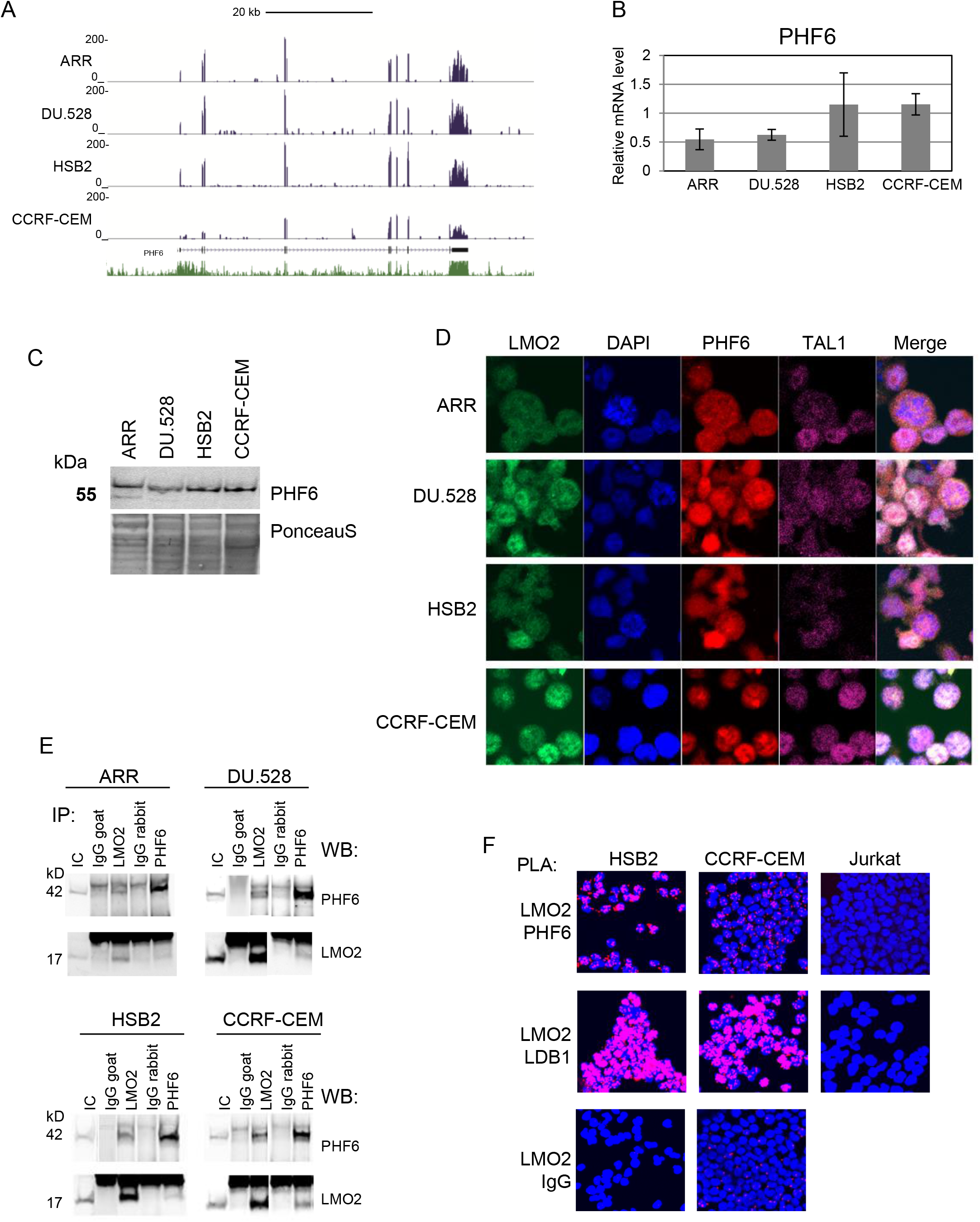
PHF6 interacts with LMO2 in T-ALL cell lines. (A) RNAseq analyses of PHF6 expression in T-ALL cell lines. Screenshot from the UCSC browser illustrating the distribution of reads over the PHF6 gene. Uniform y-axis scales were used. (B) PHF6 mRNA expression level assessed by qPCR and relative to GAPDH mRNA level. Data points are the mean of at least three independent samples measured in duplicate ± StDev. (C) Western blot showing PHF6 protein levels in T-ALL cell lines with PonceauS illustrating equal loading. (D) Confocal microscopy of immuno-fluorescently stained T-ALL cell lines, showing DNA (DAPI-blue), LMO2 (green), PHF6 (red) and TAL1 (magenta). (E) PHF6 and LMO2 co-immunoprecipitation using nuclear extracts from T-ALL cell lines. Indicated IgG were used as a control. IC: input control. Presence of PHF6 and LMO2 was visualised by western blotting (WB). (F) Proximal ligation assay (PLA) showing LMO2-PHF6 interaction in HSB2, CCRF-CEM cells. Confocal microscopy images show the nuclei visualised by DAPI (blue), while magenta staining indicates places of interaction between the indicated antibodies. LMO2-LDB1 PLA was used as a positive control, while LMO2-rabbit IgG was a negative control, as was the Jurkat T-ALL cell line as it does not express LMO2..

### PHF6 binds at the subset of LMO2/TAL1/GATA2/LDB1 target sites in T-ALL cell lines

To identify if PHF6 associates with the DNA through the LMO2/TAL/LDB/GATA complex, we performed PHF6 ChIPseq analysis. We found PHF6 peaks in all four T-ALL cell lines, with thousands of PHF6-associated regions identified in ARR, DU.528 and HSB2 cells, whereas only 250 peaks were found in CCRF-CEM cells due to low read depth from the PHF6 CCRF-CEM ChiPseq library (Figure 2A). Out of 250, 58 regions were common in all four T-ALL cell lines, while 974 were found in at least two of the cell lines (Figure 2A). Motif analyses showed that PHF6 binds to DNA fragments whose sequences were enriched for ETS, GATA, RUNX and E-box (bHLH) transcription factor binding sites but most significantly enriched was a telomeric repeat sequence (Figure 2B). Annotation of the biological processes for the genes associated with the identified PHF6 binding sites found that these DNA elements are important for haematopoietic and lymphocytic development, function and activation (Figure 2C).

**Figure 2.**
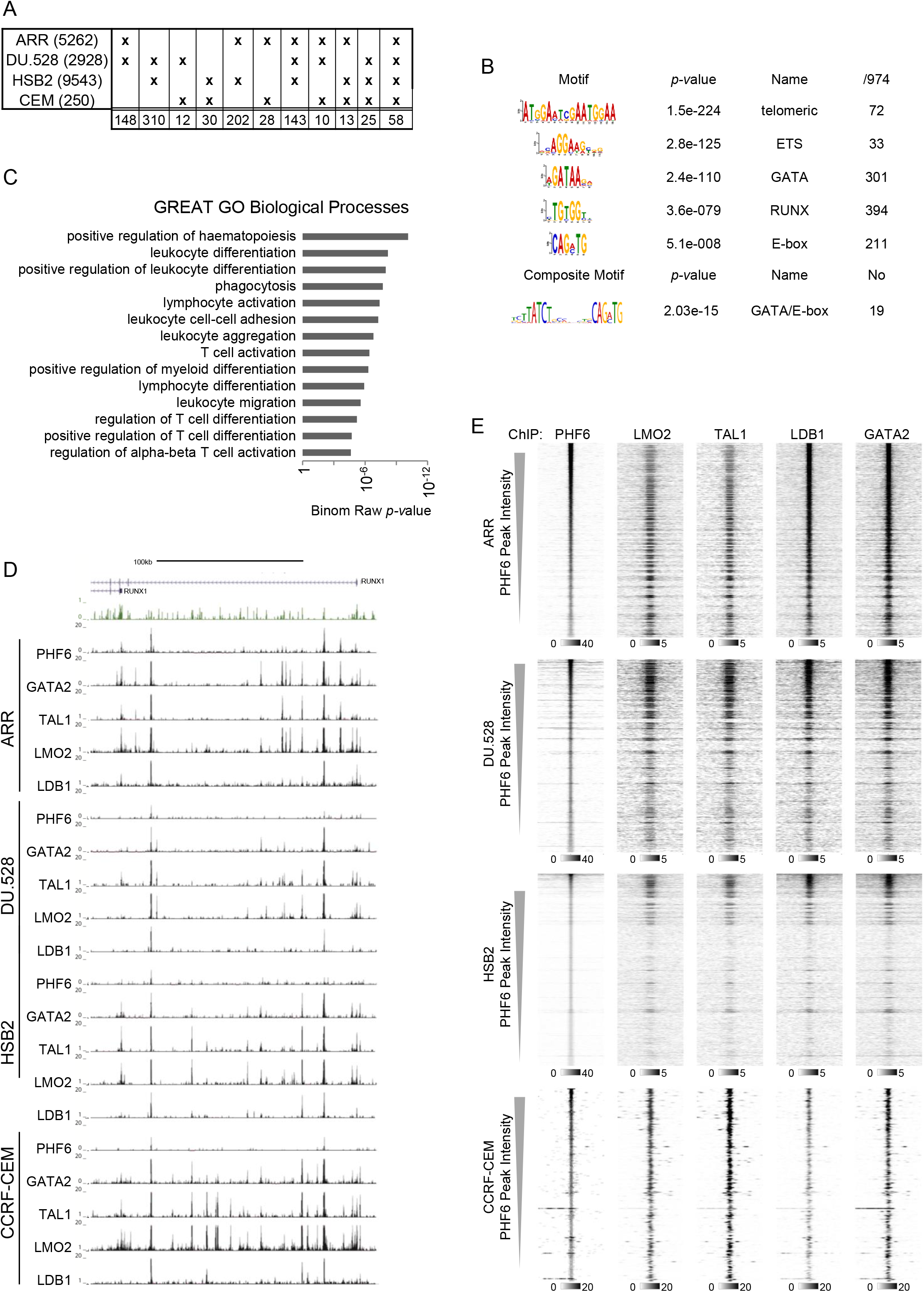
PHF6 co-localises with LMO2, TAL1, LDB1 and GATA2 at the DNA in T-ALL cell lines. (A) Table showing the overlap between PHF6 ChIPseq peaks in the four T-ALL cell lines. The first column indicates the name of the cell line and the total number of PHF6 peaks. Other columns indicate the overlaps and the number of times they occurred. (B) *De novo* motif analysis showing enriched motifs within the 974 PHF6 ChIPseq peaks that are common between at least two T-ALL cell lines. Transcription factors able to bind to the motifs, *p*-values showing the significance of the enrichment and the number of peaks found to contain the motif, are indicated. (C) GREAT gene ontology enrichment analysis for biological process performed on genomic regions of the PHF6 ChIPseq peaks. (D) Screenshot from the UCSC browser showing PHF6, GATA2, LMO2, TAL1 and LDB1 binding profiles at the *RUNX1* locus in T-ALL cell lines. Uniform y-axis scales were used for all ChIPseq tracks. (E) Heat maps showing PHF6, LMO2, TAL1, LDB1, GATA2 ChIPseq results ranked according to the intensity of PHF6 binding, depicting windows from −1 kb to +1 kb around the centre of the PHF6 peaks. The intensity of the greyscale is indicated below the heat maps.

We then performed ChIPseq for LMO2 and its binding partners TAL1, LDB1 and GATA2. Analyses of the identified binding sites confirmed that PHF6 binds DNA together with LMO2 as 40% of PHF6 peaks co-localise with identified LMO2, TAL1, LDB1 and GATA2 peaks (Figure S1). The *RUNX1* intronic enhancer is one of the DNA fragments found to be bound by PHF6-LMO2 complex in all four T-ALL cell lines and this result was confirmed in independent ChIP experiments quantified by qPCR (Figure 2D and S2). Heat maps illustrating the ChIPseq overlap with PHF6 peaks, ranked according to the PHF6-peak intensity, showed the co-localisation of PHF6 with LMO2, TAL1, LDB1 and GATA2 in all four T-ALL cell lines (Figure 2E).

### PHF6/LMO2/TAL1/LDB1/GATA2 complex has distinct function from LMO2/TAL1/LDB1/GATA2 complex in T-ALL cells

To explore the function of PHF6 in T-ALL cells we compared its binding with the LMO2 complex to that without the members of the LMO2 complex and integrated these results with our previously published RNAseq gene expression data [49]. For the benefit of concision LMO2/TAL1/LDB1/GATA2 and PHF6/LMO2/TAL1/LDB1/GATA2 binding sites will be referred to as LTLG and PHF6-LTLG peaks, respectively. First, we identified DNA segments that are occupied by PHF6 and not by any of the members of the LMO2-complex. Comparing these sites between the T-ALL cell lines found only one peak that was common for ARR, DU.528 and HSB2 cells and only 32 occupied in more than one cell line, revealing that PHF6-only sites are cell line specific (Figure 3A). The PHF6-LTLG complex was found on 58 sites in all the cell lines while an additional 165 regions were bound by three out of four LTLG factors (Figure 3B). This also means that all 58 peaks bound by PHF6 in all four T-ALL cell lines, identified during the analyses of the PHF6 ChIPseq data (Figure 2A) are bound by LMO2, TAL1, LDB1 and GATA2 (Figure 3B). These peaks associate with 161 transcriptionally active genes of which 67 (42%) were significantly differentially expressed between the T-ALL cell lines. Hierarchical clustering of the expression of these genes showed that the majority is expressed in more than one cell line albeit on a different level (Figure 3C). Functional annotation of these genes showed that they are involved in haematopoietic or lymphoid organ development, haematopoiesis, T cell differentiation and activation, transcription, chromatin organisation, positive regulation of nucleotide metabolism, intracellular transport and response to mechanical stimulus (Figure 3D). The most noticeable among these is the transcription term, as it shows that PHF6-LTLG binding sites associate with the genes encoding transcription factors such as CEBPA, RUNX1, ETS2, STAT3, STAT5A, MYBL2, JARID2, ERG and NFE2; all of which are important for the differentiation and function of the haematopoietic system.

**Figure 3.**
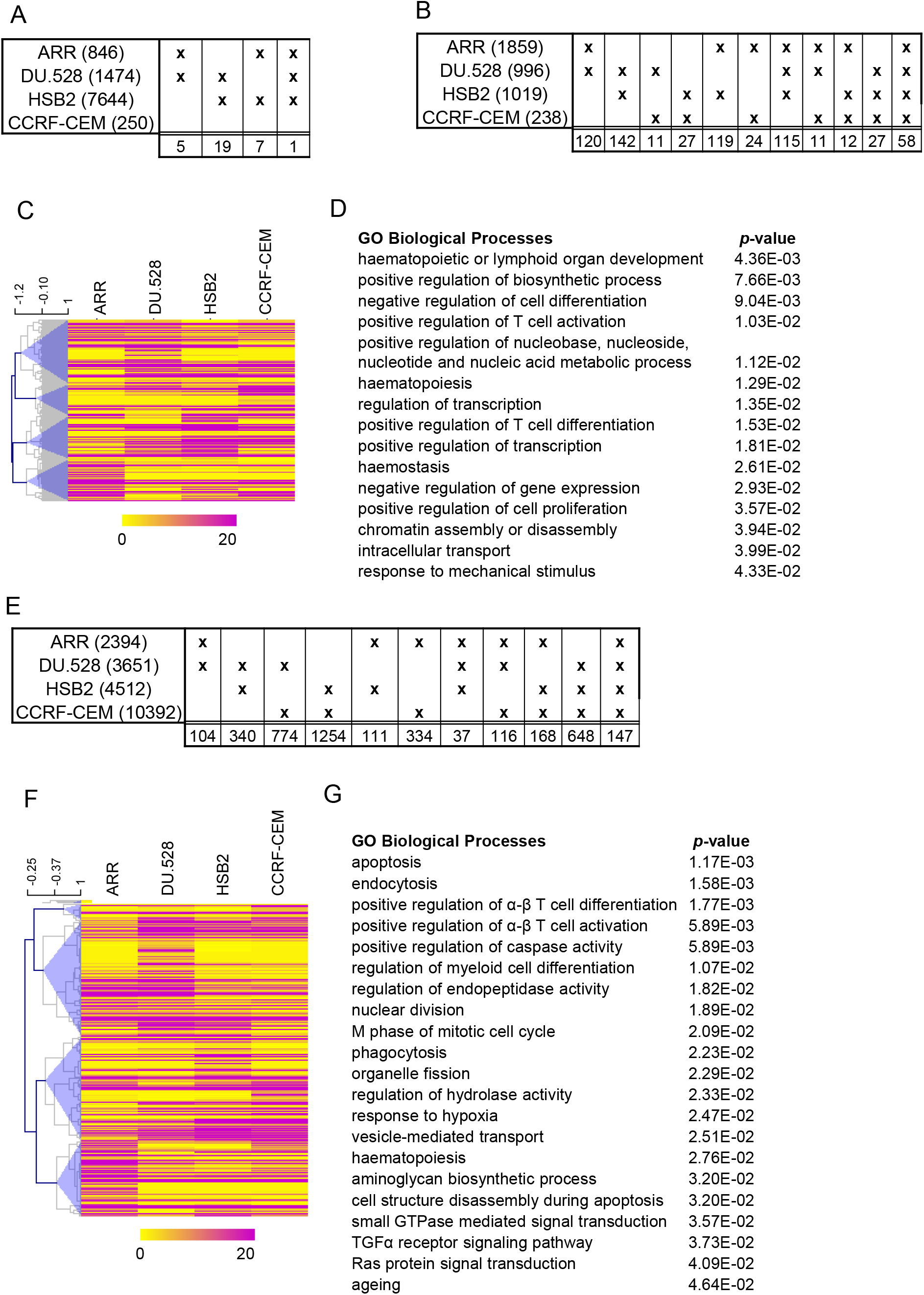
The PHF6/LMO2/TAL1/LDB1/GATA2 complex has a distinct function from the LMO2/TAL1/LDB1/GATA2 complex in T-ALL cells. (A), (B), (E) Tables showing the overlap between T-ALL cell lines of PHF6-only, PHF6-LTLG and LTLG peaks, respectively. (C), (F) Heat map showing hierarchical clustering based on the FPKM values of the nearest 5’ and 3’ gene to the PHF6-LTLG and LTLG peak, respectively. (D), (G) Gene ontology enrichment analyses for biological process was performed on the genes associated with PHF6-LTLG and LTLG peaks respectively. Terms were ordered according to their Modified Fisher Extract *p*-value and only terms with *p* < 0.05 were considered significant.

Analyses of the LTLG complex without PHF6 showed that each cell line had thousands of LTLG binding sites but only 147 DNA regions were bound by the complete complex in all four cell lines (Figure 3E). The small proportion of common LTLG peaks indicates that the localisation of the complex is more cell line specific than the PHF6-LTLG complex. Integration with the RNAseq data found that out of 275 genes that were associated with the LTLG peaks, 94 (34%) were significantly differentially expressed. Hierarchical clustering of LTLG-associated genes identified four groups of genes (Figure 3F). While the gene expression patterns of the clustering was similar to the clustering of the genes associated with PHF6-LTLG peaks, functional annotation showed that these genes have different functions, mainly relating to response to apoptosis, endocytosis, phagocytosis, nuclear division, M phase of mitosis, response to hypoxia, T-cell differentiation and activation, myeloid differentiation and haematopoiesis, as well as GTPase, TGFαR and RAS signalling (Figure 3G).

Our results showed that PHF6-binding sites that are common for all four T-ALL cell lines bound LMO2, TAL1, LDB1 and GATA2 as well. Comparison between the PHF6-LTLG and LTLG associated genes (161 and 275, respectively) identified only 6 common genes. In line with this, functional annotation showed that PHF6-LTLG and LTLG associated genes have distinct functions. PHF6-LTLG peaks were associated with genes encoding transcription factors and chromatin organisers, while LTLG peaks were found in the vicinity of genes whose products are important for signal transduction, cellular and developmental processes.

### PHF6 interacts with the LMO2-complex in Flk-1^+^ haemangioblasts and is essential for their development

The members of the LMO2-complex are essential for mesoderm specification into haemangioblasts at the onset of haematopoietic development [4, 9, 50–53]. Since LMO2 pull down followed by mass spectrometry (Supplementary file S1), identified PHF6 as an LMO2-binding partner in Flk-1^+^ haemangioblasts, we decided to establish the role of PHF6 during early haematopoietic development. Firstly, we compared the capacity of *Phf6*^-/-^, *Lmo2*^-/-^ and WT mouse ES cells to differentiate to Flk-1^+^ haemangioblasts. After four days of differentiation we found that *Phf6*^-/-^ produced embryoid bodies (EB) that were several fold larger than those generated from the WT ES cells (data not presented). Further examination of EBs revealed that *Phf6*^-/-^ cell produced 70% less Flk-1^+^ than WT and *Lmo2*^-/-^ ES cells (Figure 4A). These results show that PHF6 is required for haemangioblast development. Reduced capacity to produce Flk-1^+^ mesoderm is similar to the phenotype observed for *Ldb1*^-/-^ and *Tal1*^-/-^ ES cells [50–52]. Purification of *Phf6*^-/-^ Flk-1^+^ cells proved to be very difficult with the majority of cells not surviving the manipulations, rendering the isolation of RNA and protein extracts unproductive. However, were able to produce haemogenic endothelium (HE) from the *Phf6*^-/-^ Flk-1^+^ cells and perform fluorescent immunostainings for TIE-2 (red), CD34 (green) and KIT (magenta) (Figure 4B). Visualisation of CD34 localisation found a lack of cobblestone structure in *Phf6*^-/-^ HE that was observed as expected in the WT and Lmo2^-/-^ HE. Additionally, DNA staining (DAPI-blue) showed a number of damaged nuclei in *Phf6*^-/-^ HE. Altogether, these results show that *Phf6* is required for specification of mesoderm into haemangioblasts and haemogenic endothelium, as well as for HE genome integrity.

**Figure 4.**
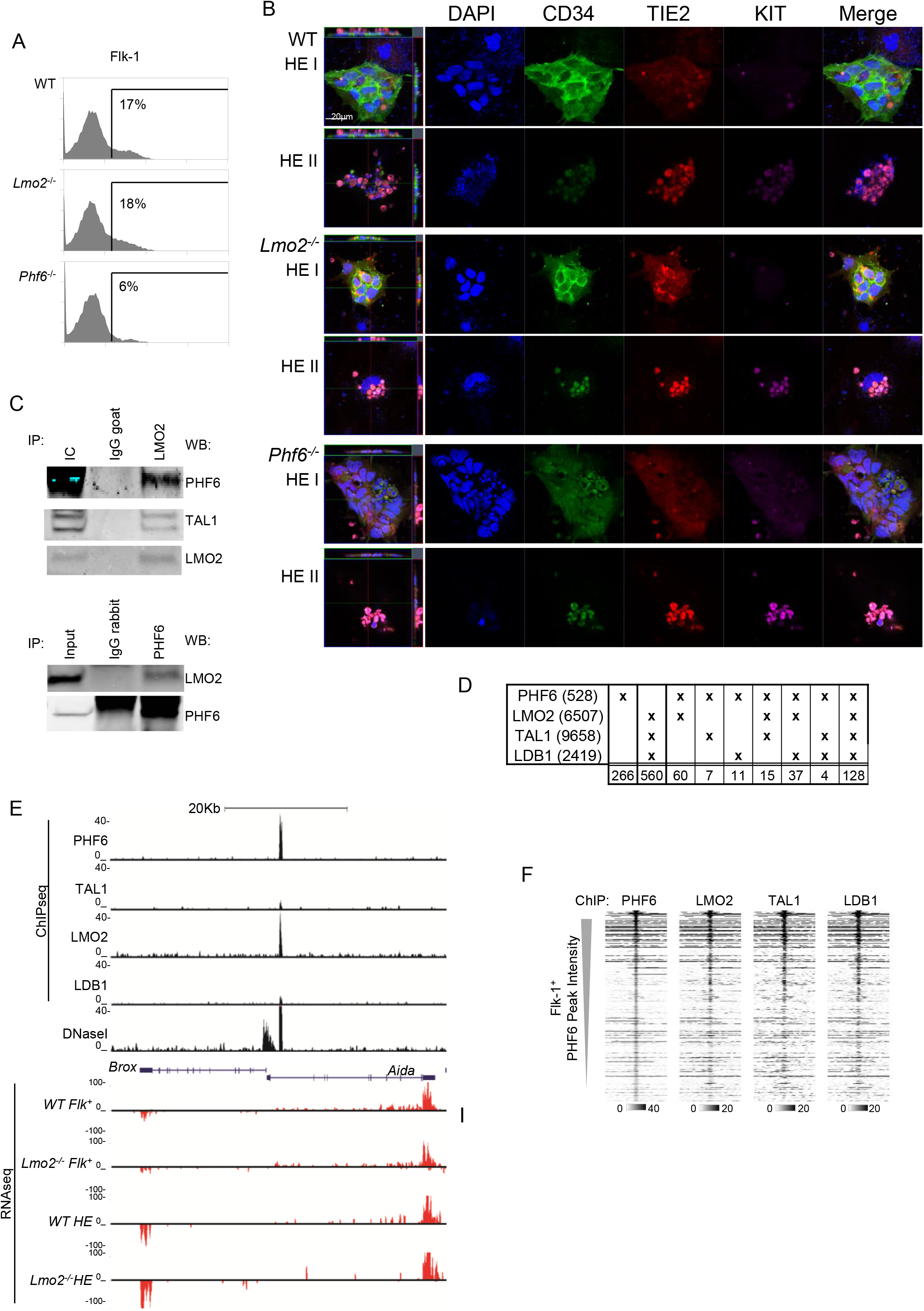
PHF6 is essential for haemangioblast development and interacts with the LMO2-complex in Flk1+ haemangioblasts. (A) WT, *Lmo2*^-/-^ and *Phf6*^-/-^ ES cells were differentiated for 3.75 days. Flk-1 surface expression was measured by flow cytometry, with prior gating for live cells using the FSC and SSC plot. Indicated percentages relate to the total number of gated live cells. (B) Confocal microscopy images of immuno-fluorescently stained HEI and HEII derived from the indicated ES cell lines. DNA (DAPI-blue), CD34 (green), TIE2 (red) and KIT (magenta) protein. First image on the left shows the collapsed Z-stack. The two lines crossing the image indicate the position along which the Z-stack projection is visualised. Projection is seen along the top and the right side of the image. Scale bar shows 20 μm. (C) PHF6 and LMO2 co-immunoprecipitation using nuclear extract from Flk-1^+^ cells. Goat IgG was used as a control. IC: input control. Results were visualised by western blotting (WB) using PHF6, LMO2 and TAL1 antibodies. (D) Table showing the overlap between PHF6, LMO2, TAL1 and LDB1 ChIPseq peaks in Flk-1^+^ cells. The first column indicates the ChIPseq and the total number of peaks. Other columns indicate the overlaps and the number of times they occurred. (E) Screenshot from the UCSC browser showing PHF6, LMO2, TAL1 and LDB1 binding profiles in WT Flk-1^+^ cells and RNAseq of WT and *Lmo2*^-/-^ Flk-1^+^ and HEI cells at the Brox and Aida locus. Uniform y-axis scales were used for all ChIPseq and RNAseq tracks. (F) Heat maps showing PHF6, LMO2, TAL1 and LDB1 ChIPseq results ranked according to the intensity of PHF6 binding, depicting windows from - 1 kb to +1 kb around the centre of the PHF6 ChIP peaks. The intensity of the greyscale is indicated below the heat maps.

LMO2 and PHF6 immunoprecipitation using nuclear extract from the Flk-1^+^ cells confirmed that the two proteins interact in these cells (Figure 4C). To assess if PHF6 binds together with the LMO2 complex we performed PHF6 ChIPseq and compared the data to the ChIPseq results we previously obtained for LMO2, TAL1 and LDB1 [4]. Analysis of the PHF6 ChIPseq data identified 526 peaks of which 24% were common for LMO2, TAL1 and LDB1, whereas 50% were shared with at least one member of the LMO2 complex (Figure 4D,E). Overlay of the LMO2, TAL1 and LDB1 ChIP data over the PHF6 binding sites, ordered by the intensity of PHF6 binding, showed that the members of the LMO2 complex bind with higher affinity to the high intensity PHF6 peaks (Figure 4F). Altogether, we showed that PHF6 is required for haemangioblast and haemogenic endothelium differentiation and that PHF6 interacts with LMO2 and binds DNA together with LMO2, TAL1 and LDB1. This, together with the observation that *Phf6*^-/-^ cells have a similar reduction in generation of haemangioblasts to *Tal1*^-/-^ and *Ldb1*^-/-^ cells, suggests that the PHF6-LMO2 interaction is required for the correct initiation of haematopoietic development.

### PHF6/LMO2/TAL1/LDB1/GATA2 complex has distinct function from LMO2/TAL1/LDB1/GATA2 complex in Flk-1^+^ cells

Further analyses of the results focused on DNA elements that were bound by PHF6 and the influence of PHF6 on the level of gene expression. *De novo* motif analysis of the PHF6 binding sites found an enrichment for telomeric repeats, but also for binding sites for NFAT, YY1, SOX8, HDX, NKX and AP-1 transcription factors (Figure 5A). Next, we wanted to explore the effect of the PHF6-DNA interaction on gene expression. As we were not able to generate RNAseq data from the *Phf6*^-/-^ Flk-1^+^ or HE cells, we used our previously published gene expression data from the WT and Lmo2^-/-^ cells [4]. Previous analyses had found 13870 transcribed genes, of which 3727 (27%) were significantly differentially expressed. Based on the genomic co-ordinates of the 528 PHF6-bound DNA elements we identified 702 associated genes that were expressed in the WT or *Lmo2*^-/-^ cells at the Flk-1^+^ or HE stage. We found that 216 (31%) genes were significantly differentially expressed. Functional annotation analyses identified that these genes are involved in the cell adhesion, embryonic development, metabolic processes, as well as mitosis and negative regulation of apoptosis (Figure 5B). With the aim to establish whether PHF6 binding is associated with a particular expression pattern we compared the expression of PHF6-associated genes to the same number of randomly selected expressed genes and performed hierarchical clustering (Figure 5C and D, respectively). The comparison between the clusters of both data sets found two clusters (C1 and C4) that were specific to the PHF6-associated set of genes and comprised of genes expressed in Flk1^+^ cells; C1 genes were suppressed in *Lmo2*^-/-^ cells compared to WT, whereas C4 genes were upregulated. Based on this analysis, the presence of PHF6 does not directly correlate to either activation or inhibition of gene expression. As we found that PHF6-bound regions are enriched for a number of transcription factor binding sites, it is possible that depending on the context of surrounding DNA-binding partners, the PHF6-containing complex has a different influence on the regulation of gene expression.

**Figure 5.**
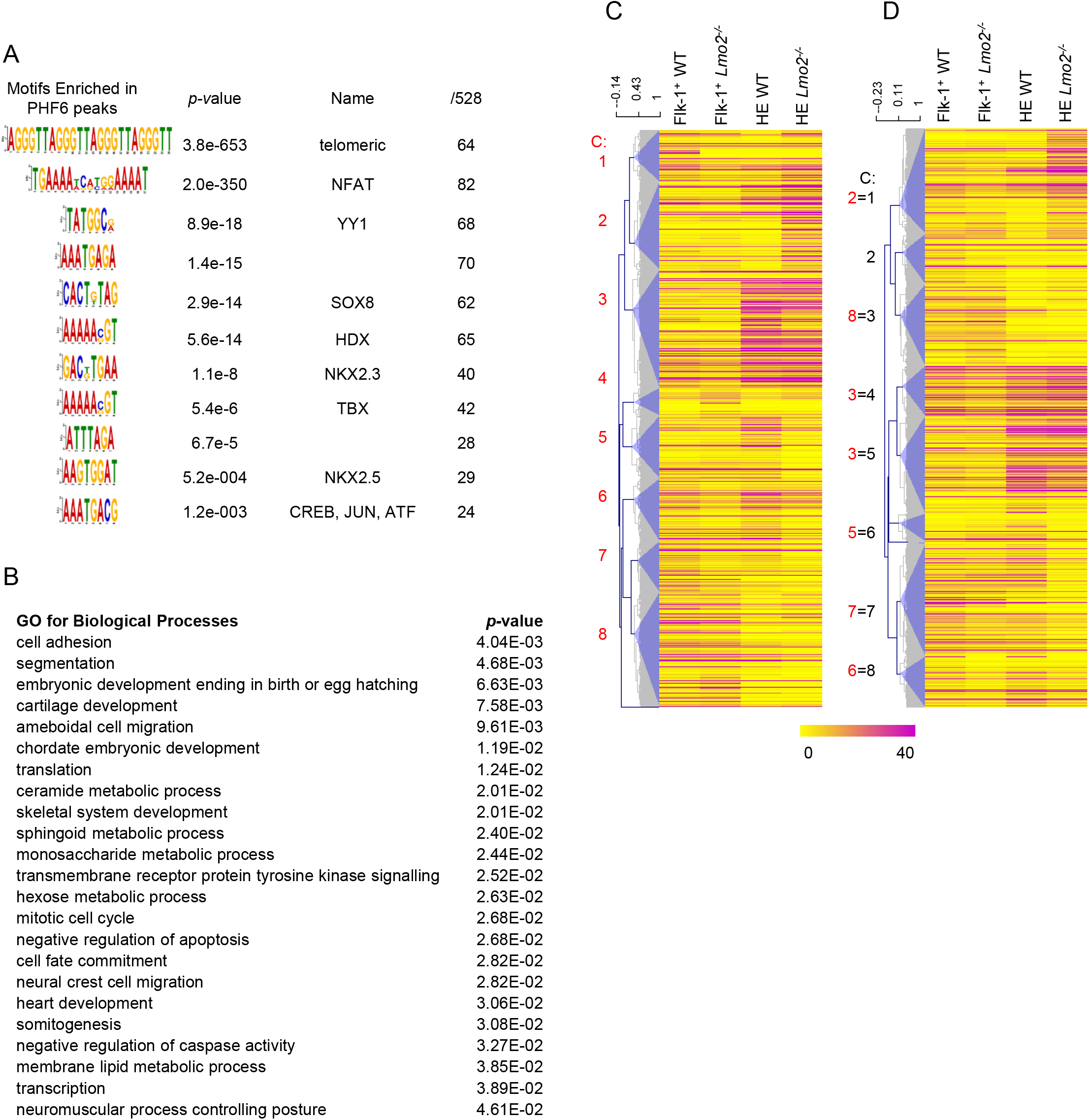
Characterisation of PHF6 binding sites in Flk-1^+^ cells. (A) *De novo* motif analysis showing enriched motifs within the 528 PHF6 ChIPseq peaks common for at least two T-ALL cell lines. Transcription factors that are predicted to bind to the motifs, *p*-values showing the significance of the enrichment and the number of peaks found to contain the motif are indicated. (B) Gene ontology enrichment analysis for biological process was performed on the nearest 5’ and 3’ expressed genes from the PHF6 peaks. Terms were ordered according to their Modified Fisher Extract *p*-value, only terms with *p* < 0.05 were considered significant. (C) Heat map illustrating hierarchical clustering of the FPKM values for the genes that are associated with the PHF6 peaks in Flk-1^+^ cells. (D) Heat map illustrating hierarchical clustering of the FPKM values for the randomly selected transcriptionally active genes in Flk-1^+^ cells.

With the aim to examine whether the PHF6-LMO2/TAL1/LDB1 (PHF6-LTL) complex has a specific role in the regulation of gene expression we analysed PHF6-LTL peaks separately from the PHF6-only peaks (Figure S2). *De novo* motif analysis found that telomeric repeat and NFAT motifs were common for both, whereas SOX8, ZNF713, HDX, E-box and NKX2.5 were enriched in Phf6-LTL peaks, and a complex motif with binding sites for the members of the AP-1 complex, E-box and RUNX1 was over-represented in PHF6-only peaks (Figure S2A,B). Functional annotation for the associated genes found significant enrichment for cell adhesion, regulation of transcription and transcription in the PHF6-LTL data, while genes associated with PHF6-only peaks were involved in cellular signalling (mTOR, FGFR, ERK, Wnt, PDGF, EGF and TGF), developmental processes (ossification, epithelial to mesenchymal transition and neuronal projection), cellular processes (Calcium-dependent cellular adhesion, endoplasmic reticulum unfolded protein response, ubiquitination and oxidation-reduction), but also ATP-dependent chromatin remodelling, negative regulation of transcription and DNA-dependent DNA synthesis (Figure S2C,D). Examination of the hierarchical clustering showed that the PHF6-LTL gene set was mostly composed of genes that were transcriptionally active in Flk-1^+^ cells and at a somewhat higher level in WT than *Lmo2*^-/-^ cells (Figure S2E). Among the genes that associated with PHF6-only peaks (Figure S2F), those with higher expression in WT HE were more abundant.

Next, we examined the genes that are in the vicinity of LMO2/TAL1/LDB1 peaks that do not overlap with PHF6 binding by hierarchical clustering and found that two main clusters (Figure S2G, C1 and 2) were characterised by genes with higher expression in WT Flk-1^+^ compared to *Lmo2*^-/-^ Flk-1^+^ cells. The expression of these genes was further enhanced from WT Flk-1^+^ to HE stage, demonstrating that the LMO2/TAL1/LDB1 complex without PHF6 is required for activating gene expression at the onset of haematopoiesis (Figure S2G). Gene ontology analysis revealed that these genes are important for transcription, translation, cell cycle, nucleotide and fatty acid metabolism, as well as for a number of developmental processes (Figure S2H).

Similar to our observations in the T-ALL cell lines, the results from the Flk-1^+^ cells show that PHF6 interacts with LMO2. PHF6 is required at the onset of haematopoietic development when mesoderm is specified to develop into haemangioblasts and haemogenic endothelium thereafter. The PHF6-LTL, PHF6-only and LTL-only complexes associate with distinct gene sets with diverse gene ontologies. Moreover, we were able to discriminate between the LTL-complex with or without PHF6. The PHF6-LTL-complex was important for activating gene expression in the Flk-1^+^ stage, while the LTL complex associated more with genes that were expressed at a higher level in WT Flk-1^+^ than in *Lmo2*^-/-^ cells and whose expression was upregulated at the HE stage.

### PHF6 interacts with the LMO2 complex during myeloid differentiation

As PHF6 mutations are also found in AML patient samples, we decided to establish if PHF6 interacts with the LMO2 complex during myeloid differentiation. For this we used the PUER cell line, which are common myeloid progenitors derived from *Spi1* (encoding PU.1 transcription factor) knock out mice. PUER cells are engineered to express a fusion of PU.1 with the ligand binding domain of oestrogen receptor, which results in active nuclear PU.1 upon the addition of 4-hydroxytamoxifen (OHT) to the medium. Induction of PU.1 initiates differentiation of PUER cells to macrophages. PLA assays confirmed the interaction of PHF6 with LMO2 (Figure 6A).

**Figure 6.**
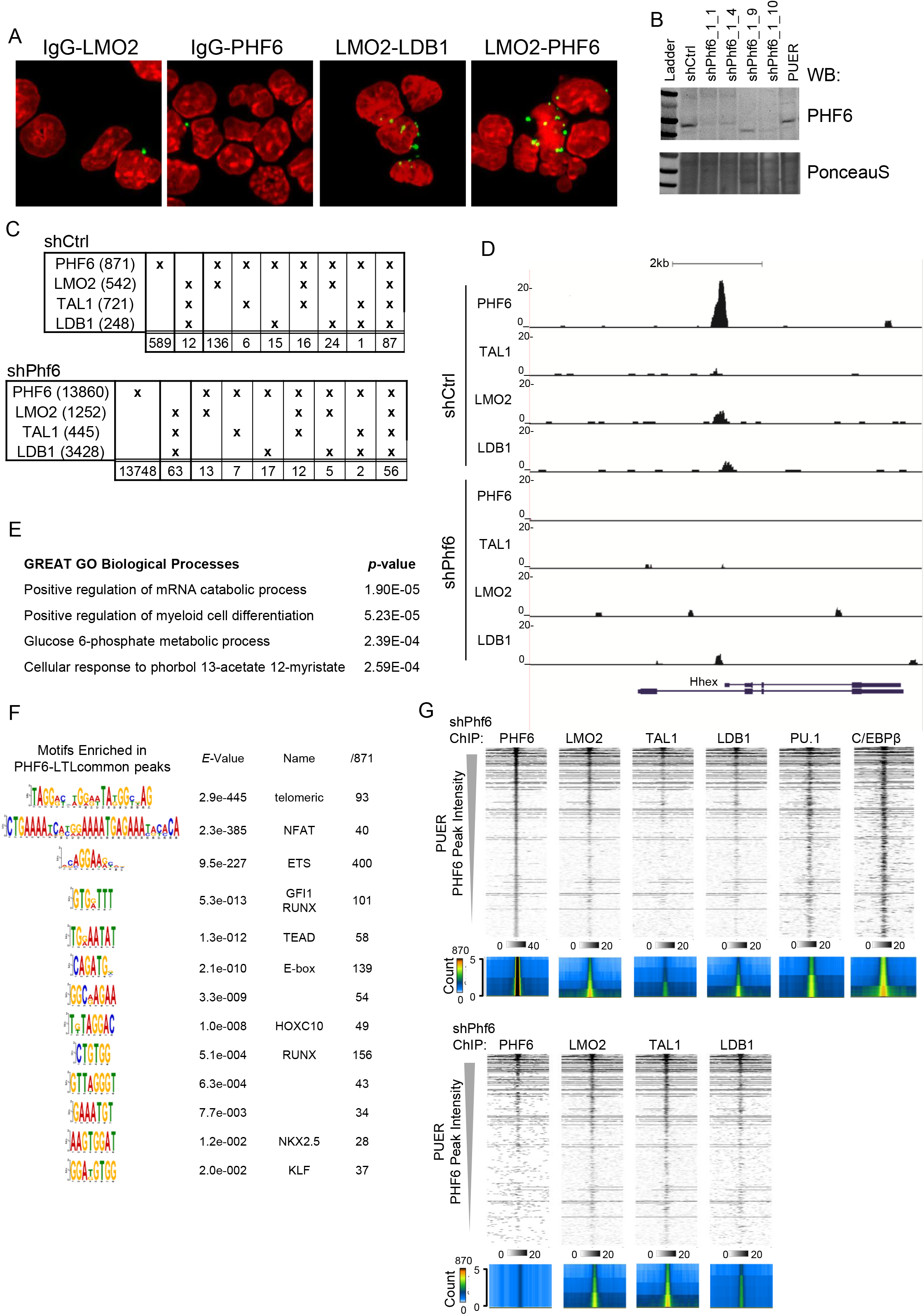
PHF6 interacts with the LMO2 complex during myeloid differentiation. (A) LMO2-PHF6 interaction in mouse myeloid common progenitor cell line (PUER) was detected in proximal ligation assay (PLA). LMO2-LDB1 PLA was used as a positive control while LMO2-abbit IgG and PHF6-goat IgG were negative controls. Confocal microscopy images show nuclei visualised by DAPI DNA stain (red) while the green dots show the places of the interaction between the indicated antibodies. (B) PHF6 protein levels shown by western blot from the PUER shCtrl and shPHF6_1 single cell clones. Equal loading was visualised PonceauS staining. (C) Table showing the number of overlapping peaks between PHF6, LMO2, TAL1 and LDB1 ChIPseq in PUER shCtrl and shPHF6 cells, respectively. Total number of peaks is indicated on the left, while the numbers at the bottom row indicate overlapping peaks. (D) Screenshot from the UCSC browser showing PHF6, LMO2, TAL1 and LDB1 binding profiles at the Hhex locus in shCtrl and shPHF6 PUER cells. Uniform y-axis scales were used. (E) GREAT gene ontology enrichment analysis for biological process performed on genomic regions associated with the PHF6 ChIPseq peaks. (F) *De novo* motif analysis showing enriched motifs within the 871 PHF6 ChIPseq peaks detected in shCtrl PUER cells. Transcription factors that are predicted to bind the motifs, *p*-values showing the significance of the enrichment and the number of peaks containing the motif are indicated. (G) Heat maps showing PHF6, LMO2, TAL1, LDB1, PU,1 and C/EBPβ ChIPseq results from shCtrl PUER cells (top), and PHF6, LMO2, TAL1, LDB1, ChIPseq results from shPHF6 PUER cells, both ranked according to PHF6 peak intensity in shCtrl cells. Windows from –1 kb to +1 kb around the centre of the PHF6 peaks. The intensity of the greyscale is indicated below the heat maps. Below each matrix an average profile shows the distribution of the ChIP signal throughout the matrix.

To characterise the importance of PHF6 and its interaction with LMO2 in myeloid cells we generated shPHF6 PUER cell lines with reduced PHF6 levels. We tested six different shPhf6 sequences and examined their efficiency to reduce *Phf6* expression. Efficiency was assayed by expressing PHF6 together with GFP from a poly-cistronic mRNA and measuring the level of GFP expression after introduction of each shPhf6 construct, using shCtrl and shGFP as negative and positive controls respectively (Figure S3A). Only shPHF6_2 was not effective in supressing PHF6-GFP expression. shPHF6_1, _5 and _6 were used to generate shPHF6 PUER cell lines, but we were only able to generate a stable cell line with shPHF6_1, while PUER transduced with shPHF6_5 and shPHF6_6 did not grow, As shPHF6_5 and shPHF6_6 showed the greatest knockdown efficiency this indicates the importance of PHF6 for survival of the common myeloid progenitors. PUER shPhf6_1 was used for the generation of single cell clones, with clone shPHF6_1_1 showing a clear reduction of endogenous PHF6 without additional bands in the western blot, which was therefore subsequently used for further experiments (Figure 6B). To test the capacity of shPhf6 cells to differentiate into macrophages we induced their differentiation and assessed the expression levels of the macrophage differentiation markers CD11b and F4/80. Compared to the control, shPhf6 exhibited normal upregulation of CD11b, a known direct PU.1 target gene, but had reduced F4/80 levels, showing that PHF6, in addition to being required for the survival of myeloid progenitors, also has a role in myeloid differentiation (Figure S3B).

To find out whether the importance of PHF6 in myeloid cells is due to its interaction with the LMO2 complex we performed ChIPseq experiments for PHF6, LMO2, TAL1 and LDB1 in shCtrl and shPhf6 cells (Figure 6C,D). We identified 870 PHF6 binding sites, 87 (10%) of which were shared with LMO2, TAL1 and LDB1 (PHF6-LTL) and 282 (30%) were bound by PHF6 and at least one of the LMO2 complex members. The *Spi1* enhancer is one of the DNA fragments found to be bound by the PHF6-LMO2 complex in PUER cells and this result was confirmed in independent ChIP experiments quantified by qPCR (Figure S3C). These results confirmed that in myeloid cells PHF6 interacted with the LMO2 complex (Figure 6C). In shPhf6, the number of PHF6-occupied DNA elements was 13,860 and a similar increase in the number of binding sites was observed for LMO2 and LDB1 while the number of TAL1 binding sites was reduced. The PHF6-LTL complex was also found in shPhf6 cells but only on 57 genomic locations, despite the large number of peaks. Analyses of the gene ontology associated with the PHF6 binding sites in shCtrl cells, using the GREAT database found enrichment for mRNA catabolism, myeloid differentiation, glucose metabolism and cellular response to phorbol 12-myristate 13-acetate (PMA), which activates PKC signaling and induces myeloid differentiation and activation [54–57] (Figure 6E). A search for enriched DNA motifs within the PHF6 binding regions revealed that, in addition to telomeric repeat and NFAT binding sites, ETS DNA-binding motifs had the highest enrichment as GGAAG was detected in 400 out 872 examined sequences. Binding sites for RUNX, GFI1/GFI1b, E-box, NKX2.5 and KLF were found as well (Figure 6F). Overlap of the ChIPseq data ordered according to the intensity of PHF6 binding in the shCtrl and shPhf6 cells illustrated that the intensity of LMO2, TAL1 and LDB1 binding correlated with PHF6 binding (Figure 6G). As the ETS motif was found in a great number of PHF6 peaks, we used previously published PU.1 ChIPseq data to find whether PHF6 and PU.1 co-localise at the same DNA regions [58]. We used PU.1 ChIPseq acquired from PUER cells prior to the induction of differentiation. Overlap of these two data sets showed that PU.1 co-localised with PHF6 (Figure 6G). This was confirmed when we focused the analysis on PU.1 peaks which were ordered by the intensity of PU.1 binding and then overlaid with our data which showed the highest correlation with PHF6 and LMO2 followed by LDB1 and TAL1 (Figure S3D). A previously reported finding that CEBPβ binds the same elements as PU.1, was confirmed for PHF6-PU.1 bound DNA elements (Figure 6G). In shPhf6 cells we observed less intense PHF6 signal, as expected, but also increased TAL1 binding, suggesting that PHF6 may limit the interaction of TAL1 with LMO2 in myeloid progenitors (Figure 6G).

For further analysis PHF6 peaks were separated into PHF6-only peaks and peaks that overlap with LMO2 and/or TAL1 and/or LDB1 (PHF6-L/T/L). Functional annotation for the genes associated with PHF6-only peaks found that these DNA elements are important for phagocytosis, mRNA catabolism, protein glycosylation and myeloid cell differentiation, whereas the PHF6-L/T/L peaks correlated to stimulation of the GTPase activity (Figure S3E). Performing *de novo* motif enrichment analysis for Phf6-L/T/L peaks showed that NFAT, ETS, E-box, IRX3, GFI1, ZSCAN4 and PAX5 binding sites were more frequent while Phf6-only peaks had overrepresented E-box-ETS composite motif, E-box and RUNX motifs (Figure S3F). Altogether we conclude that PHF6 interacts with LMO2 in myeloid cells and that the reduced PHF6 level leads to increased TAL1 DNA binding as well as an increase in the number of LTL-peaks, suggesting that PHF6 limits the availability of the LMO2/TAL1 complex for binding to the DNA.

### Phf6 knock down perturbs myeloid gene expression

To determine the importance of PHF6 for myeloid differentiation we performed RNAseq for shCtrl and shPhf6 cells prior to differentiation (D0), as well as 24h (D1) and 72h (D3) after induction of differentiation. Analysis of the results identified 281 upregulated and 249 downregulated genes in pairwise comparison of shPhf6 cells with treatment-matched shCtrl cells (Figure 7A). Interestingly, we observed a two–and–a-half times higher number of upregulated genes in shPhf6 (52) than downregulated (20) prior to differentiation, which indicates that overall PHF6 suppresses gene expression. As expected, among the differentially expressed gens we found supressed PHF6 expression in shPHF6 cells at D0 and D1, whereas *Adgre1*, encoding the F4/80 protein, was downregulated at D1 and D3 (Figure S4A,B), in line with the flow cytometry data (Figure S3A). Gene ontology analysis of the differentially expressed genes found that their function is important for a) the activity of transcription factors, b) haematopoietic, myeloid, lymphoid and adipose development c) fatty acid biosynthesis, inhibition of glucose transport and d) inflammation, locomotion, wound healing and immune response (Figure 7B).

**Figure 7.**
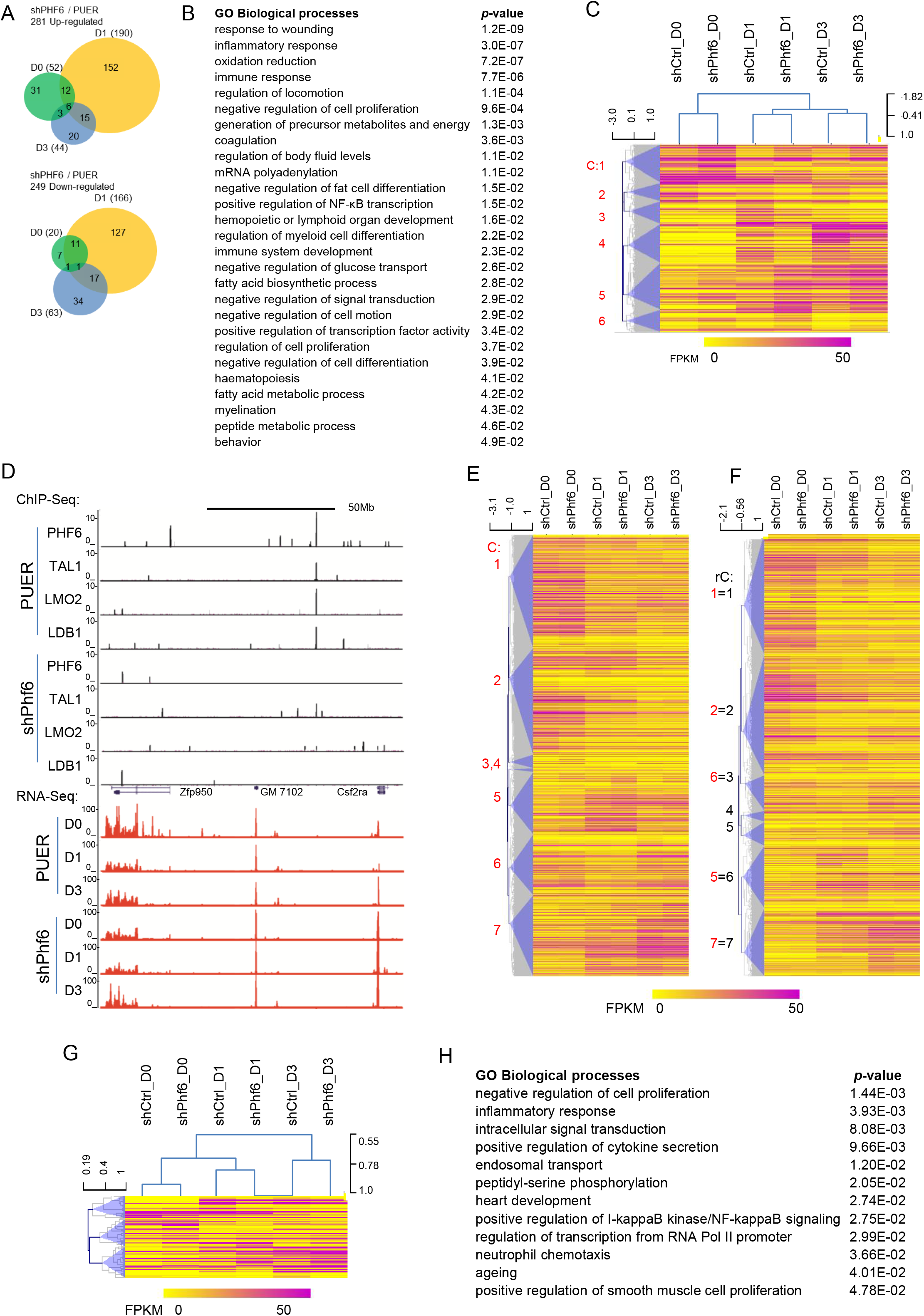
*Phf6* knock down perturbs myeloid gene expression. (A) Three-way Venn diagram showing the numbers of significantly differentially expressed genes between shCtrl and shPHF6 PUER cells at D0, D1 and D3 of differentiation. Shown are the total number of up or down regulated genes for each comparison and their overlap. Quantitation and analyses of RNAseq data are based on biological duplicate samples. (B) Gene ontology enrichment analysis for biological process for significantly differentially expressed genes between the shCtrl and shPHF6 PUER cells. Terms were ordered according to their Modified Fisher Extract *p*-value, only terms with *p* < 0.05 were considered significant. (C) Heat map showing hierarchical clustering of all differentially expressed genes. Scale bar represents colour index for the FPKM values. Six clusters are identified and numbered 1 to 6. (D) UCSC genome browser screenshot showing the distribution of reads across *Zfp950* and *Csf2Ra* from PHF6, LMO2, TAL1 and LDB1 ChIPseq and RNAseq tracks for shCtrl and shPHF6 PUER cells. Uniform y-axis scales were used for all ChIPseq and RNAseq tracks. (E) Heat map illustrating hierarchical clustering of FPKM values for the genes that are associated with the PHF6 peaks in shCtrl and shPHF6 PUER cells. Seven clusters are identified (C1-7) (F) Heat map illustrating hierarchical clustering of the FPKM values for an equal number of randomly selected transcriptionally active genes in shCtrl and shPHF6 PUER cells. Seven clusters are identified (rC1-7) and comparable clusters to those in (F) are indicated. (G) Heat map illustrating hierarchical clustering of FPKM values for significantly differentially expressed genes between shCtrl and shPHF6 PUER cells that associate with PHF6 peaks. (H) Gene ontology enrichment analysis for biological process using the significantly differentially expressed genes between the shCtrl and shPHF6 PUER cells that associate with the PHF6 peaks. Terms were ordered according to their Modified Fisher Extract *p*-value, only terms with *p* < 0.05 were considered significant.

Hierarchical clustering of the differentially expressed genes identified six clusters (C1-C6, Figure 7C). C1, C5 and C6 were comprised of genes that were higher in shPhf6 cells, whereas C2, C3 and C4 were lower. In addition to the effect of *Phf6* knock down, differentiation had a major effect on gene expression. Genes organised in C1 and C2 were downregulated during differentiation, C3 genes were transiently expressed at D1, while genes in C4 and C5 were upregulated from D0 to D3. Based on the hierarchical clustering we noticed that, in addition to a two-and-half fold higher number of upregulated genes in shPHF6 cells at D0, the majority of genes that were expressed at D0 were expressed at a higher level in PHF6 knock down cells (C1, C5 and C6). This strongly suggests that PHF6 contributes to the suppression of gene expression. Among the genes that were upregulated at D0 we found transcription factors *Egr1, Nfkbia* and *Fos*, which shows that PHF6 could exert its suppressive function directly or indirectly by supressing the expression level of other transcription factors.

With the aim to determine whether PHF6 directly regulates the expression of the differentially expressed genes we integrated RNAseq and ChIPseq analyses and identified 1752 genes in the vicinity of PHF6 binding sites. Many of the PHF6 peaks were associated with multiple genes and some of these were significantly differential expressed between the shCtrl and shPHF6, such as *Csf2ra* and *Zfp950* (Figure 7D). Interestingly, the genes associated with PHF6/LMO2/TAL1/LDB1 peaks were largely identical to those associated with PHF6-only peaks, rendering analysis of independent groups redundant. Therefore we focused on exploring the expression of all PHF6-associated genes. To establish whether PHF6 binding is associated with a particular developmental expression pattern or levels, we again generated a set of randomly-selected expressed genes and subjected both sets to hierarchical clustering. Visualisation showed that the majority of PHF6-associated genes were expressed at higher levels than the equivalent clusters in the random gene set (*e.g*. compare C1, C5, C6, C7 to rC1, rC6, rC3 and rC7 respectively) (Figure 7E, F), which was confirmed by pairwise comparison of the two gene sets for D0, D1 and D3 (Figure S4C). Therefore we conclude that PHF6-bound regions are associated with genes whose level of expression is relatively higher.

Among the PHF6-associated genes, 94 were significantly differentially expressed between shPhf6 and the control, in comparison to 52 found in the random gene set, indicating that PHF6 peaks associate more commonly with the genes whose expression changed as a result of the PHF6 knock down. Clustering these genes according to the level of expression showed that the majority had higher expression in the shPHF6 cells at D0 and their expression can decrease (C2) or increase (C3) during differentiation, while a smaller number started being expressed at D1 and were higher in shCtrl (C1) (Figure 7G). Gene ontology showed that these genes were involved in negative regulation of cell proliferation, inflammatory response, endosomal transport, heart development, positive regulation of I-κB kinase/NF-κB signalling, regulation of transcription from RNA Pol II promoter and ageing (Figure 7H). Among the transcription factors that were associated with PHF6 binding we found *Fos*, *Nfkbia* and *Egr1*, which were significantly upregulated in shPHF6 cells prior to differentiation (D0). This indicates that PHF6 is required for restricting the expression levels of transcription factors that are required for myeloid differentiation and activation.

### PHF6 and LMO2 are essential for genomic stability

Thus far we showed that PHF6 interacts with LMO2 and together they bind to DNA in T-ALL cells, Flk-1^+^ haemangioblasts and myeloid progenitors. The PHF6-LMO2 complex associated with different genes that are important for the function of each of the explored systems. In order to distinguish which effects of PHF6 knock down are due to its interaction with LMO2 we generated *Lmo2* knock down PUER cells (shLmo2). Three different shLmo2 were made and all were effective (Figure S5A). Comparing the appearance of the cells, we noticed that while shCtrl cells had a typical myeloid appearance with doughnut like nuclei, many shLmo2 and shPhf6 cells were greater in size with large lobular nuclei, which is indicative of dysregulated replication/cell division (Figure 8A). Mitotic chromosome spreads found that the majority of shCtrl had a normal diploid genome (40 chromosomes), whereas in shPhf6 and shLmo2 cell cultures, approximately one third had between 40 and 100 chromosomes with another third having more than 100 chromosomes per cell (Figure 8B,C). The morphology of the chromosomes in some of the aberrant mitosis was that of pulverised chromosomes. Based on this we conclude that reduction of LMO2 or PHF6 leads to a loss of chromosome stability.

**Figure 8.**
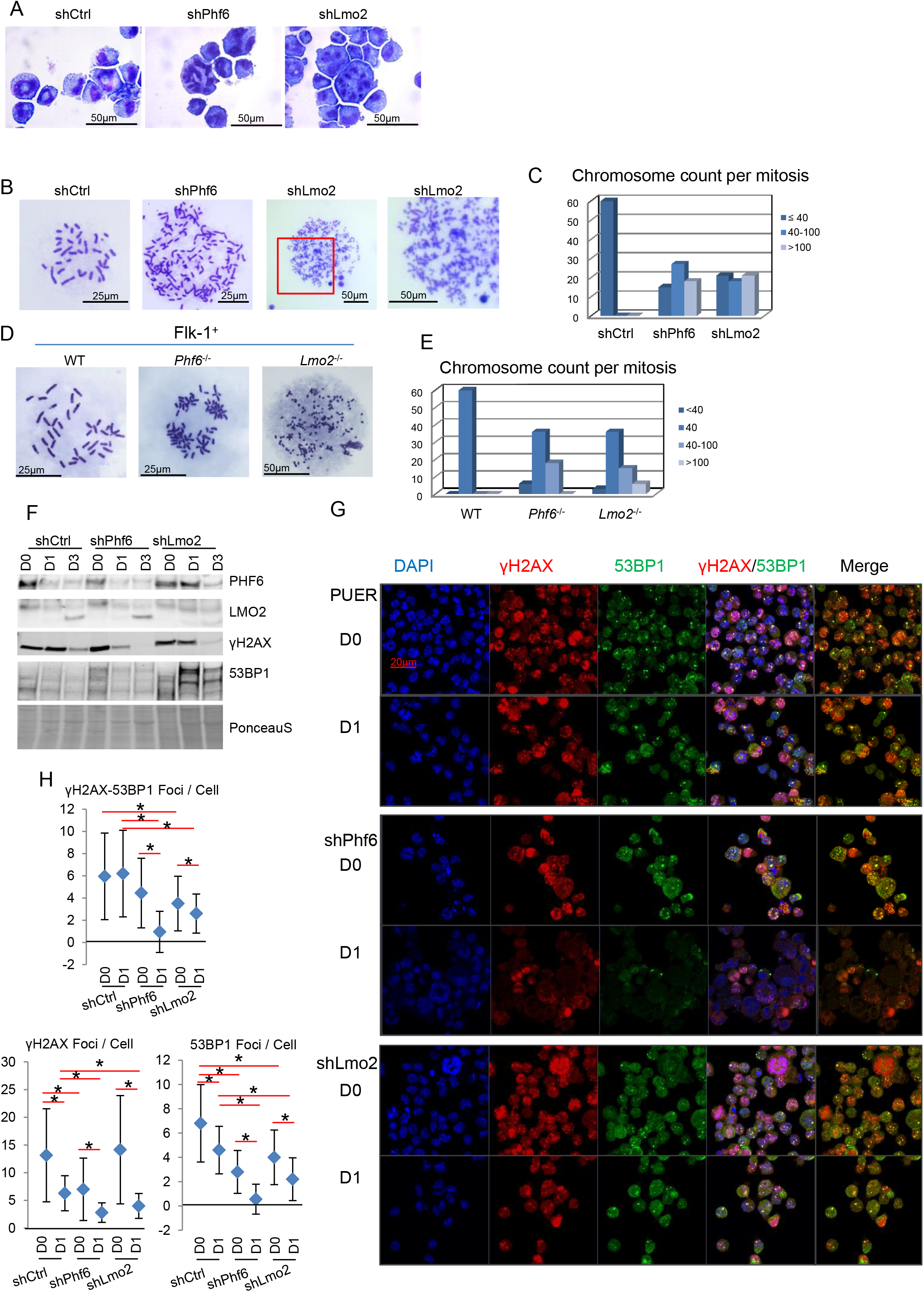
PHF6 and LMO2 are essential for genomic stability. (A) Cellular morphology shown by histochemical staining of shCtrl, shPhf6 and shLmo2 PUER cells. (B) Representative mitotic chromosome spreads of shCtrl, shPhf6 and shLmo2 cells. Larger magnification of the shLmo2 panel is shown on the right. (C) Bar graph showing the number of mitoses containing up to 40, 40-100 and more than 100 chromosomes for the indicated PUER cell lines. (D) Representative mitotic chromosome spreads of the WT, *Phf6*^-/-^ and *Lmo2*^-/-^ Flk-1^+^ cells. (E) Bar graph showing the number of mitoses containing <40, 40, 40-100 and >100 chromosomes for indicated ES cell-derived Flk-1^+^ cells. (F) Western blot showing the protein levels of PHF6, LMO2, γH2AX and 53BP1 in PUER cell lines at the indicated stages of differentiation. PonceauS illustrates equal loading. (G) Confocal microscopy images of immuno-fluorescent staining showing the distribution and co-localisation of the γH2AX (red) and 53BP1 (green) protein in PUER cells lines prior and 1 day after the initiation of differentiation. DNA was stained with DAPI (blue). Merged images are shown on the right. Scale bar shows 5 μm. (H) Graphs showing the average number of γH2AX and or 53BP1 foci per cell. Twenty cells were assessed and the average number ± standard deviation was presented. One-tail heteroscedastic T-test was used for calculation of the *p*-values. Relevant comparisons with significant differences are shown.

Similar observations were made for *Lmo2*^-/-^ and *Phf6*^-/-^ Flk-1^+^ cells, where we found that approximately half of the cells had a normal number of chromosomes, about 10% were hypoploid and the remaining 40% had an increased number of chromosomes, with some of the *Lmo2*^-/-^ displaying fragmented chromosomes (Figure 8D,E). These results show that the PHF6-LMO2 complex is required for maintaining chromosome integrity.

The observed increase in the number of chromosomes indicates that the cells are endomitotic, where they go through mitosis without cell division. A possible reason for this is a reduced ability to repair DNA damage that is left unresolved. This is even more likely as PHF6 depletion is reported to be correlated with increased γH2AX levels [21, 30, 33]. To investigate the importance of LMO2 and PHF6 for DNA repair we assessed the level of γH2AX and 53BP1 in shCtrl, shPhf6 and shLmo2 cells prior to and D1 and D3 after the initiation of differentiation. We found that in comparison to the control, shPhf6 have increased levels of γH2AX and 53BP1 at D0, which is followed by a reduction of γH2AX at D1 and undetectable levels at D3. The shLmo2 cells had similar γH2AX levels to the control, but clearly higher 53BP1 levels at all time points. Normally, γH2AX is recruited to damaged DNA where it nucleates with 53BP1 and the DNA repair machinery. Therefore the cellular distribution and co-localisation are important indicators of the capacity of cells to repair DNA damage. To find out how changes in the total γH2AX and 53BP1 levels related to their intracellular distribution and co-localisation, we used immuno-fluorescent staining. Examination of the microscopic images and quantification of γH2AX/53BP1 foci revealed that in comparison to shCtrl, shLmo2 cells had reduced number of γH2AX/53BP1 foci at D0 and D1, while shPhf6 cells only showed a statistically significant decrease at D1 (Figure 8G,H,S5B). Closer examination of γH2AX and 53BP1 separately showed that both shPhf6 and shLmo2 showed reduced numbers of 53BP1 foci at D0 and D1. Even though shPhf6 cells did not have a significantly lower number of γH2AX/53BP1 foci at D0, they had lower numbers of γH2AX at this time point, indicating that in these cells 53BP1 efficiently co-localised with γH2AX. However, the reduced number of γH2AX foci showed that PHF6 is needed for the recruitment of γH2AX to DNA lesions. Compared to the control, shLmo2 cells showed a similar number of γH2AX foci at D0. Therefore the observed reduction in 53BP1 foci at D0, directly translated to the reduction of γH2AX/53BP1 foci, showing that LMO2 has a function in associating 53BP1 to the places of γH2AX accumulation (Figure 8G,H,S5B). Together, these findings illustrate that LMO2 and PHF6 are important for the DNA damage response, with PHF6 being required for γH2AX accumulation and LMO2 for recruiting 53BP1 to the γH2AX foci.

Altogether, our results show that PHF6 interacts with LMO2 as a part of the TAL1, LMO2, LDB1 complex in T-ALL, mES cell-derived Flk-1^+^ haemangioblasts and in myeloid progenitors. *Phf6*^-/-^ ES cells inefficiently generated Flk-1^+^ cells, which further differentiated to haemogenic endothelium, lacking CD34 expression and the normal cobblestone-like organisation. We show that the PHF6-LMO2 complex binds to the DNA and in myeloid cells these binding sites are important for down-regulating the level of gene expression that are required for myeloid differentiation and function. In addition to the function in transcription we show that PHF6 and LMO2 are important for the maintenance of the normal number of chromosomes and this could in part be due to their role in recruiting the DNA damage–repair machinery. We show that PHF6 and LMO2 are required for maintaining normal levels of γH2AX and 53BP1, where PHF6 is important for γH2AX accumulation and LMO2 has a role in recruiting 53BP1 to γH2AX foci.

## Discussion

LMO2 is a transcriptional mediator that through its interaction with transcription factors TAL1, LYL and FOG/GATA binds to DNA, whereas its contacts with LDB1, which can trimerise, facilitate the interaction between distant DNA elements [1–3, 6, 7]. Other proteins were shown to interact with the LMO2/LDB1 complex such as SP1, ELK1, NFATC1, LEF1 and SSBP [59–61]. In this paper, we identified PHF6 as a novel LMO2-interacting partner. Analysis of the ChIPseq data showed that PHF6 occupied different DNA elements in different cells and a proportion of these were bound by LMO2 as well. However, the overlays show that the majority of PHF6 peaks are enriched for LMO2 binding. This difference is most likely due to the low intensity of the interactions or as a consequence of the insufficient depth of ChIPseq data. Due to the nature of the LMO2 complex and its importance for mediating long-distance interactions, it is possible that ChIPseq experiments also enrich for DNA elements that are indirectly bound by the PHF6-LMO2 complex.

TAL1 and LDB1 ChIPseq data identified that these two factors also bound the same DNA fragments as PHF6 and LMO2. We showed that this interaction occurs during normal haematopoiesis, as well as in leukaemia. *In vitro* ES differentiation found that *Phf6*^-/-^ ES cells have a reduced ability to generate Flk-1^+^ haemangioblasts, which was not the case for *Lmo2*^-/-^ ES cells (herein and [4]). In this aspect, *Phf6*^-/-^ differentiation shows similarity with the *Ldb1*^-/-^ and *Tal1*^-/-^ phenotypes [50, 62]. In the CMP cell line PUER, in addition to observing the PHF6-LMO2 interaction, we found that a reduction in PHF6 was followed by an increase in TAL1 binding. This could be due to increased TAL1 protein levels upon PHF6 depletion or as a consequence of PHF6 competition with TAL1 for interaction with LMO2. Even though we show that PHF6 interacts with the LMO2/LDB1/TAL1 complex, PHF6 could favour a particular variant of this complex, as TAL1 can be replaced by LYL1 or HEB, LMO2 by LMO1, and LDB1 by LDB2.

In addition to PHF6, LMO2, TAL1 and LDB1, we report the association of additional factors such as GATA2 in T-ALL and PU.1, C/EBPβ in PUER cells. This shows that the PHF6-LMO2 complex interacts with lineage-specific partners. As LMO2 nucleates TAL1, GATA and LDB1, the contribution of PHF6 could be to facilitate the interactions with additional tissue-specific factors.

Several publications reported different roles for PHF6. PHF6 was shown to interact with the NuRD (Nucleosome Remodelling Deacetylase) complex [29, 31, 63]. The NuRD complex, which contains CHD3/4 (Chromodomain Helicase DNA Binding Protein, also known as Mi-2a and b), RBBP (retinoblastoma binding protein) and HDAC1/2 (histone deacetylase), facilitates ATP-dependent chromatin remodelling and histone deacetylase activities, leading to suppression of gene expression. We confirmed the HDAC1 interaction with PHF6 and LMO2 by co-immnoprecipitation in T-ALL cell lines (data not shown) but ChIPseq experiments for HDAC1 showed very little binding and we did not find enrichment on DNA elements that were bound by PHF6 and LMO2. This could be due to the limited capability of the antibodies that were used in the experiments. Therefore our current findings do not exclude the possibility of an interaction of the PHF6-LMO2 complex with the NuRD complex.

PHF6 was also shown to be important for the expression of ribosomal RNA as it was found to locate to the nucleolus and, through the interaction with UBF, binds to the regulatory elements of the rDNA genes [30, 64]. Our PHF6 and LDB1 ChIPseq experiments indicated enrichment for rDNA, but this was not restricted to the rDNA regulatory elements that are located upstream and downstream of transcribed rDNA. As the rDNA is composed of repeated sequences that encode ribosomal RNA and are located on five different chromosomes in humans, observed enrichments could be an artefact due to the nature of the repeats and the inability to precisely assemble the rDNA genomic sequence. The results of the PHF6 and LDB1 ChIPseq prompted us to explore further. We tried to reproduce the published data by using the same primers for measuring the enrichment of the PHF6 binding but we found that the PCR employing these primers produced multiple different DNA molecules rendering the quantification of enrichment impossible. We were able to design primers uniquely recognising the rDNA regulatory regions but in these experiments we were not able to find enrichment for PHF6. Therefore, our results failed to confirm that PHF6 binds to the regulatory regions of rDNA but it is possible that this is a cell-type specific function of PHF6 and not one it assumes in haematopoietic cells. Additionally, differences in the procedures for chromatin preparation, cell treatments and antibodies recognising different domains of PHF6 are possible reasons for the discrepancies in the findings.

Finally, we compared our data to previously published PHF6 ChIPseq data [65, 66]. Firstly, we used data deposited to GEO but due to the type and / or the quality of the submitted data, further processing using the standard workflow of the data did not produce the alignment track or peaks. Therefore we could only compare published screenshots of the PHF6 alignments to our data and we did not find PHF6 binding to any of the reported regions. Furthermore, we have used Jurkat cell line as well as the cell lines used in this publication in combination with the same antibody as reported by Meacham et al., as well as the one that we used in this study. Unfortunately, we could not confirm the previously reported findings. A different publication reported that TNFα induces PHF6/NFkB co-localisation in a BCR-ABL^+^ B-ALL cell line [35]. Comparison to this study was hampered by the quality of the PHF6 ChIPseq data and unfortunately not possible. These difficulties can be attributed to the quality of the antibodies that are available as we found a lot of variations between different PHF6 antibodies provided by a wide range of suppliers. Moreover, we observed several different forms of PHF6 that could have different cell-type specific interacting partners, which could explain the involvement of PHF6 in facilitating various cellular processes.

One consistent finding reported by several groups and herein is that reduced PHF6 availability leads to increased γH2AX levels [21, 30, 33]. Our observation that PHF6 interacts with LMO2 and that both are important for genomic integrity indicates that the PHF6-LMO2 interaction is required for chromosome integrity. Chromosome fragmentation and pulverisation cannot be maintained for a long time as it leads to cell death. This is potentially overcome by the emergence of clones that carry additional mutations, minimising the effects of the PHF6 and LMO2 depletion. We were aware of this possibility as we met serious difficulties generating and expanding single cell clones. Additionally, during the normal culture of the successfully generated clone we observed instabilities that have prompted us to control for the efficiency of PHF6 knock down on a regular basis and with the experiments reported in this publication. This level of vigilance is necessary in order to secure consistent results that address the function of PHF6 and its binding partners. Taking this in consideration would aid the production of reproducible findings and would help us develop a better understanding of PHF6 and its contribution to malignancies.

## Acknowledgments

We would like to thank Dr. A.W. Langerak, (Erasmus Medical Centre, Rotterdam, NL) for the provision of ARR and DU.528 cell lines and Prof P.N. Cockerill (University of Birmingham, UK) for HSB2 and CCRF-CEM. We would like to acknowledge BlueBEAR High Performance Computing (HPC) service for supporting the analyses of the genome-wide data. SB was supported by King Saud University and the Saudi Arabia Cultural Bureau. This work was further supported by Bloodwise, through a Bennett Fellowship to M.H. [11002], and the University of Birmingham.

## Authorship Contributions

Original Concept M.H. and V.S.S., Project Planning and Supervision, V.S.S., M.H.; Experimentation, Data Acquisition, Processing and Analyses V.S.S., S.B., I.D. and M.H.; Genome-wide data acquisition and Bioinformatical analyses V.S.S., S.B. and M.H.; Statistical analyses V.S.S. and S.B.; Mass-spectrometry data acquisition and analyses V.S.S., S.B. and D.G.W.; Figures V.S.S and S.B.; Writing of the Manuscript, V.S.S., S.B., D.G.W. and M.H.

## Notes

https://www.ncbi.nlm.nih.gov/geo/query/acc.cgi?acc=GSE154675

### Competing Interest Statement

The authors have declared no competing interest.

