## supplemental legends and figures for "PHF6 Interacts with LMO2 During Normal Haematopoiesis and in Leukaemia and Regulates Gene Expression and Genome Integrity"

Legends for Supplementary Figures

Figure S1. (A) Table showing the number of PHF6, LMO2, TAL1, LDB1 and GATA2 overlapping peaks in each of the T-ALL cell lines. Overlap between at least two of the ChIPseq peaks are coloured according to the number with low numbers in red and highest in green. (B) The same as A but for top 1000 peaks with highest intensity per ChIPseq sample. (C) Enrichment for the TAL1, LMO2, LDB1, GATA2 and PHF6 ChIP at the *Runx1* intronic enhancer and the negative control region Chr.18, relative to the input control. Data points are the mean of at least three independent samples measured in duplicate ± StDev.

Figure S2. (A), (B) *De novo* motif analysis showing enriched motifs in Flk-1$^+$ cells within the PHF6-LTL and PHF6-only peaks, respectively. Transcription factors that are predicted to bind the enriched motifs, *p*-values showing the significance of the enrichment and the number of peaks found to contain the motif are indicated. (C), (D), (H) Gene ontology enrichment analysis for biological process was performed on the nearest 5' and 3' expressed gene from the PHF6-LTL, PHF6-only and LTL-peaks, respectively. Terms were ordered according to their Modified Fisher Extract *p*-value, only terms with $p < 0.05$ were considered significant. (E), (F), (G) Hierarchical clustering of FPKM values for the genes associated with respectively the PHF6-LTL, PHF6-only and LTL peaks in Flk-1$^+$.

Figure S3. PUER shPhf6. (A) Efficacy of shRNA sequences targeting *Phf6* compared to shGFP and shCtrl, as assessed by flow cytometry of the GFP protein levels in PlatE cells. Percentage of GFP$^+$ cells is indicted. Experiment was performed three independent times and representative results are shown. (B) Surface expression levels of the myeloid differentiation markers CD11b and F4/80 by shCtrl and shPHF6_1 PUER cells prior to and during differentiation (D0, 1, 2 and 3) as measured by flow cytometry. Indicated percentages relate to the total number of live cells, based on the FSC and SSC plot. Experiment was performed three independent times and representative results are shown. (C) Relative enrichment for the PHF6, LMO2, TAL1 and LDB1 PHF6 ChIP in shCtrl and shPHF6 PUER cells at the *Spi1* (encoding PU.1) upstream enhancer and the negative control region Chr.2, relative to the input control. Data points are the mean of at least three independent samples measured in duplicate ± StDev. (D) Heat maps showing PHF6, LMO2, TAL1, LDB1, PU,1 and C/EBPβ ChIPseq results from the shCtrl PUER cells ranked according to the intensity of the PU.1 ChIPseq peaks. Window from –1 kb to +1 kb around the centre of the PU.1 peak is used and the maximum intensity of the heat map greyscale is indicated below the heat map. Below each matrix an average profile shows the distribution of the ChIP signal throughout the matrix. (E) Gene ontology enrichment analysis for biological process was performed on the nearest 5’ and 3’ expressed genes from the PHF6-LTL and PHF6-only and LTL-peaks, respectively. Terms were ordered according to their Modified Fisher Extract *p*-value, only terms with $p < 0.05$ were considered significant. (F) *De novo* motif analysis showing enriched motifs within the 282 PHF6-LTL and 589 PHF6-only ChIPseq peaks detected in shCtrl PUER cells. Transcription factors that are predicted to bind the identified motifs, *p*-values showing the significance of the enrichment and the number of peaks found to contain the motif are indicated.

Figure S4. RNAseq analysis of shCtrl and shPhf6 PUER cells. (A), (B) UCSC genome browser screenshots from a representative set of D0, D1 and D3 shCtrl and shPHF6 RNAseq tracks showing the distribution of reads across the *Phf6* and *Agdre1* (encoding F4/80) gene, respectively. (C) Table showing average FPKM and standard deviation values for the genes found 5’ and 3’ from the PHF6 ChIPseq peak or the same number of randomly chosen expressed genes. Expression was calculated for differentiation samples at D0, D1 and D3. *p*-value established based on one-tailed paired T-test.

Figure S5. (A) Efficacy of shRNA sequences targeting *Lmo2* compared to shGFP and shCtrl, as assessed by flow cytometry of the GFP protein levels in PlatE cells. Percentage of $GFP^+$ cells is indicated. Experiment was performed three independent times and representative results are shown. (B) Immunostaining of shCtrl and shPhf6 PUER cells during differentiation  Confocal microscopy images of immuno-fluorescent staining showing the distribution and co-localisation of γH2AX (red) and 53BP1 (green) protein in PUER cell lines prior to and 1 day after the initiation of differentiation. DNA was stained with DAPI (blue). Merged images are shown on the right. Scale bar shows 20 μm.

Figure S1

A

| | | | | | | | | | | | | | | | | | | | | | |
|---|---|---|---|---|---|---|---|---|---|---|---|---|---|---|---|---|---|---|---|---|---|
| PHF6 | x | | | | | x | x | x | x | x | x | x | x | x | x | x | x | x | x | | x |
| LMO2 | | x | | | | x | | | | x | x | x | | | | x | x | x | | x | x |
| TAL1 | | | x | | | | x | | | x | | | x | x | | x | x | | x | x | x |
| LDB1 | | | | x | | | | x | | | x | | x | | x | x | | x | x | x | x |
| GATA2 | | | | | x | | | | x | | | x | | x | x | | x | x | x | x | x |

| | | | | | | | | | | | | | | | | | | | | | |
|---|---|---|---|---|---|---|---|---|---|---|---|---|---|---|---|---|---|---|---|---|---|
| ARR | 846 | 10088 | 11326 | 6727 | 36864 | 39 | 18 | 322 | 218 | 32 | 11 | 81 | 0 | 12 | 1447 | 12 | 421 | 66 | 8 | 2338 | 1775 |
| DU.528 | 1474 | 8592 | 7350 | 5800 | 2310 | 11 | 20 | 43 | 19 | 35 | 3 | 6 | 0 | 8 | 154 | 40 | 132 | 7 | 4 | 3627 | 986 |
| HSB2 | 7644 | 13815 | 5087 | 3752 | 9258 | 53 | 27 | 79 | 120 | 55 | 5 | 30 | 2 | 15 | 279 | 15 | 219 | 18 | 0 | 4405 | 994 |
| CCRF-CEM | 4 | 24905 | 9616 | 11604 | 8499 | 0 | 0 | 0 | 0 | 1 | 0 | 0 | 0 | 0 | 11 | 0 | 2 | 1 | 0 | 10240 | 213 |

B

| | | | | | | | | | | | | | | | | | | | | | |
|---|---|---|---|---|---|---|---|---|---|---|---|---|---|---|---|---|---|---|---|---|---|
| PHF6 | x | | | | | x | x | x | x | x | x | x | x | x | x | x | x | x | x | | x |
| LMO2 | | x | | | | x | | | | x | x | x | | | | x | x | x | | x | x |
| TAL1 | | | x | | | | x | | | x | | | x | x | | x | x | | x | x | x |
| LDB1 | | | | x | | | | x | | | x | | x | | x | x | | x | x | x | x |
| GATA2 | | | | | x | | | | x | | | x | | x | x | | x | x | x | x | x |

| | | | | | | | | | | | | | | | | | | | | | |
|---|---|---|---|---|---|---|---|---|---|---|---|---|---|---|---|---|---|---|---|---|---|
| ARR | 171 | 111 | 132 | 192 | 373 | 5 | 4 | 401 | 8 | 42 | 1 | 1 | 2 | 1 | 0 | 19 | 82 | 1 | 3 | 57 | 185 |
| DU528 | 428 | 63 | 0 | 233 | 274 | 9 | 15 | 68 | 34 | 33 | 6 | 2 | 6 | 3 | 37 | 54 | 45 | 5 | 2 | 148 | 254 |
| HSB2 | 511 | 70 | 72 | 226 | 261 | 5 | 3 | 101 | 16 | 26 | 4 | 0 | 2 | 3 | 38 | 32 | 45 | 0 | 2 | 170 | 212 |
| CCRF-CEM | 36 | 82 | 66 | 196 | 287 | 2 | 4 | 12 | 2 | 7 | 1 | 0 | 5 | 1 | 11 | 13 | 4 | 2 | 5 | 332 | 145 |

C ChIP qPCR in T-ALL cell lines

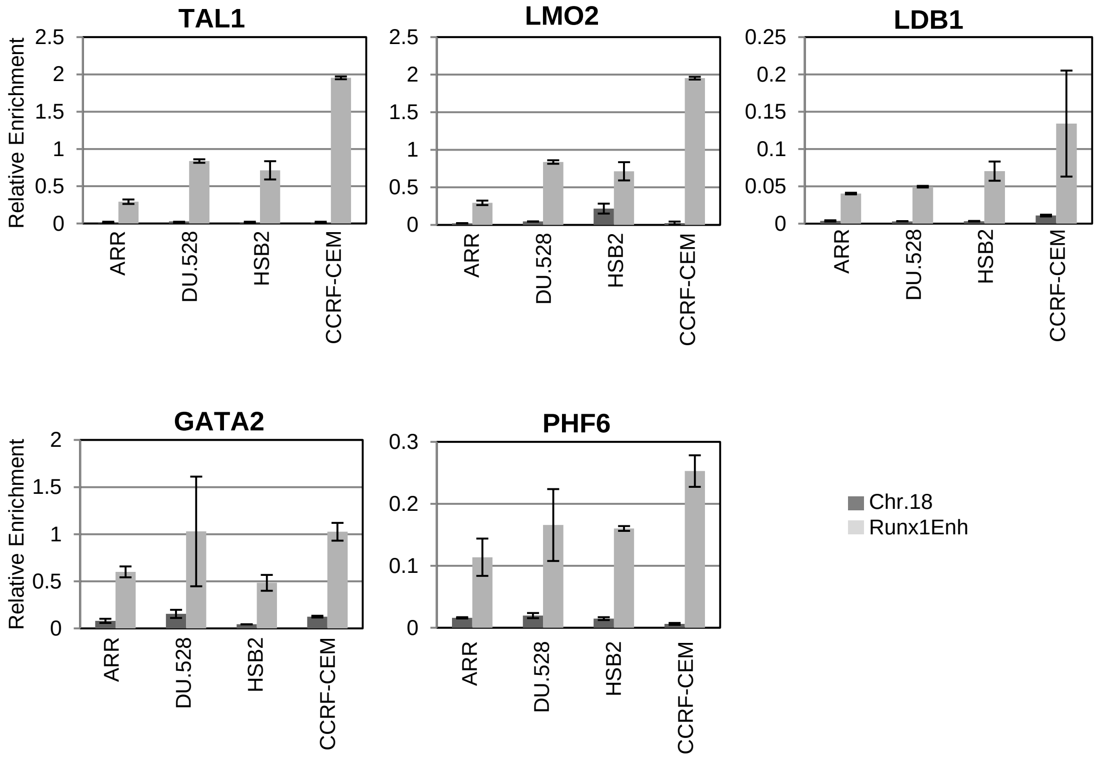

### Figure S2

A

| Motifs Enriched in PHF6-LTLcommon peaks | *p-value* | Name | /128 |
|---|---|---|---|
| 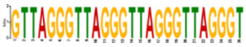 | 9.1e-455 | telomeric | 41 |
| 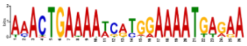 | 7.4e-285 | NFAT | 40 |
| 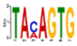 | 5.0e-007 | SOX8 | 37 |
| 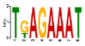 | 5.1e-007 | ZNF713 | 40 |
| 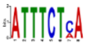 | 5.1e-007 | HDX | 33 |
| 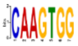 | 3.5e-002 | E-box NKX2.5 | 11 |

B

| Motifs Enriched in PHF6-only peaks | *p-Value* | Name | /257 |
|---|---|---|---|
| 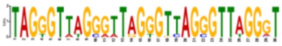 | 5.0e-177 | telomeric | 20 |
| 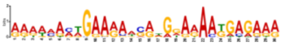 | 3.6e-116 | NFAT | 34 |
| 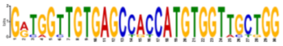 | 5.0e-007 | JUNDM2 E-box RUNX | 9 |

C

| GO Biological Processes | p-value |
|---|---|
| cell adhesion | 5.34E-03 |
| regulation of transcription | 1.53E-02 |
| transcription | 1.74E-02 |

D

| GO Biological Processes | p-value |
|---|---|
| negative regulation of ossification | 1.13E-03 |
| negative regulation of TOR signaling | 2.97E-03 |
| positive regulation of epithelial to mesenchymal transition | 6.94E-03 |
| calcium-dependent cell-cell adhesion | 8.45E-03 |
| neurotransmitter biosynthetic process | 1.30E-02 |
| fibroblast growth factor receptor signaling pathway | 1.58E-02 |
| regulation of ventricular cardiac muscle cell action potential | 1.65E-02 |
| negative regulation of ERK1 and ERK2 cascade | 1.66E-02 |
| negative regulation of extrinsic apoptotic signaling | 1.70E-02 |
| ATP-dependent chromatin remodeling | 1.82E-02 |
| oxidation-reduction process | 1.98E-02 |
| regulation of catalytic activity | 2.02E-02 |
| endoplasmic reticulum unfolded protein response | 2.28E-02 |
| Wnt signaling pathway | 2.33E-02 |
| cell division | 2.36E-02 |
| regulation of circadian rhythm | 2.96E-02 |
| T cell mediated cytotoxicity | 3.36E-02 |
| positive regulation of organ growth | 3.36E-02 |
| negative regulation of fibroblast proliferation | 3.49E-02 |
| positive regulation of neuron projection development | 3.71E-02 |
| cellular response to hydrogen peroxide | 3.95E-02 |
| response to virus | 4.14E-02 |
| negative regulation of transcription from RNA Pol II promoter | 4.24E-02 |
| filopodium assembly | 4.39E-02 |
| positive regulation of protein ubiquitination | 4.64E-02 |
| platelet-derived growth factor receptor signaling pathway | 4.70E-02 |
| response to ischemia | 4.70E-02 |
| cellular response to epidermal growth factor stimulus | 4.70E-02 |
| cellular response to transforming growth factor beta stimulus | 4.88E-02 |
| DNA-dependent DNA replication | 4.95E-02 |

E

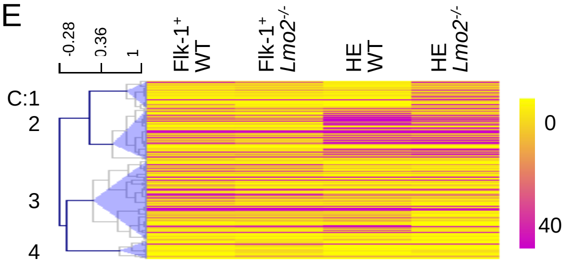


F

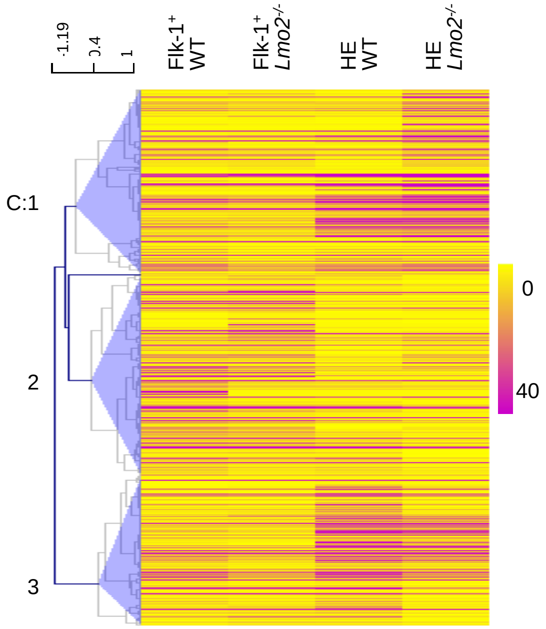


G

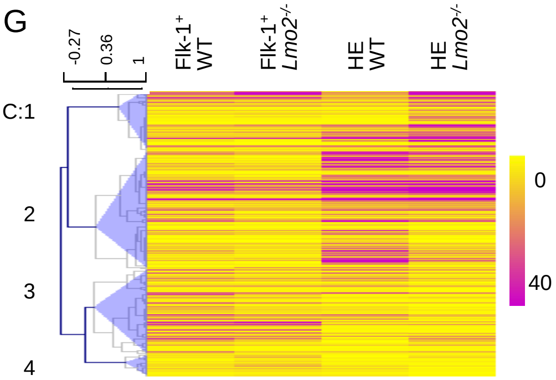


H

| GO Biological Processes | p-value |
|---|---|
| positive regulation of transcription | 2.19E-04 |
| positive regulation of nucleobase, nucleoside, nucleotide and nucleic acid metabolic process | 4.81E-04 |
| positive regulation of nitrogen compound metabolism | 6.72E-04 |
| positive regulation of transcription from RNA Pol II promoter | 1.42E-03 |
| phosphorus metabolic process | 4.17E-03 |
| cell morphogenesis | 4.50E-03 |
| epithelial differentiation in prostate gland development | 1.05E-02 |
| phosphorylation | 1.39E-02 |
| cell morphogenesis involved in differentiation | 1.49E-02 |
| enzyme linked receptor protein signaling pathway | 1.84E-02 |
| branching involved in salivary gland morphogenesis | 1.97E-02 |
| skeletal system development | 2.31E-02 |
| fatty acid metabolic process | 2.48E-02 |
| axon guidance | 3.14E-02 |
| cellular response to hormone stimulus | 3.34E-02 |
| regulation of translation | 3.35E-02 |
| posttranscriptional regulation of gene expression | 3.44E-02 |
| cell cycle | 3.53E-02 |
| cellular cation homeostasis | 3.79E-02 |

### Figure S3

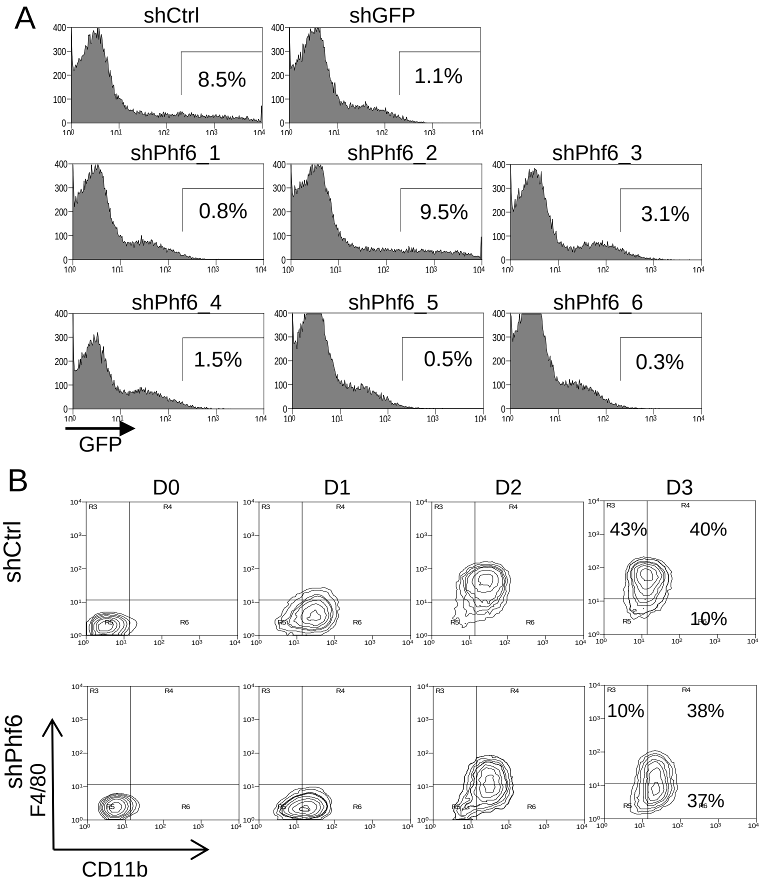


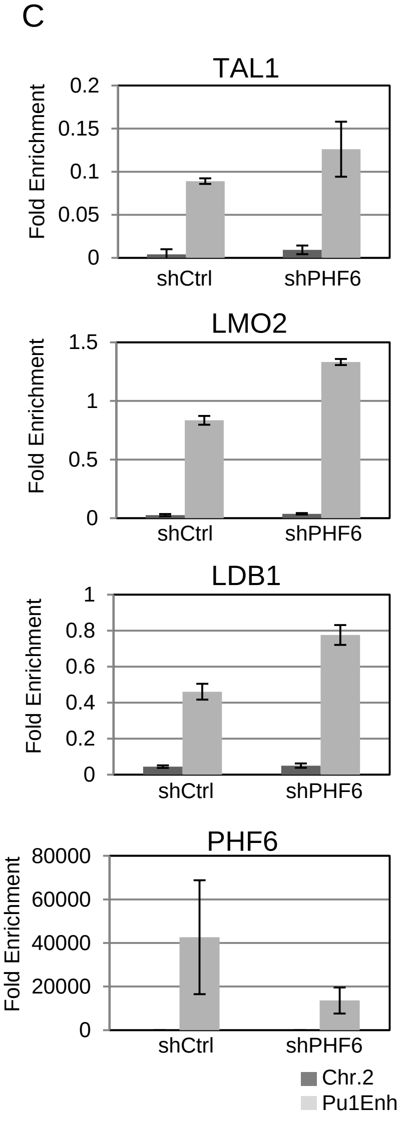


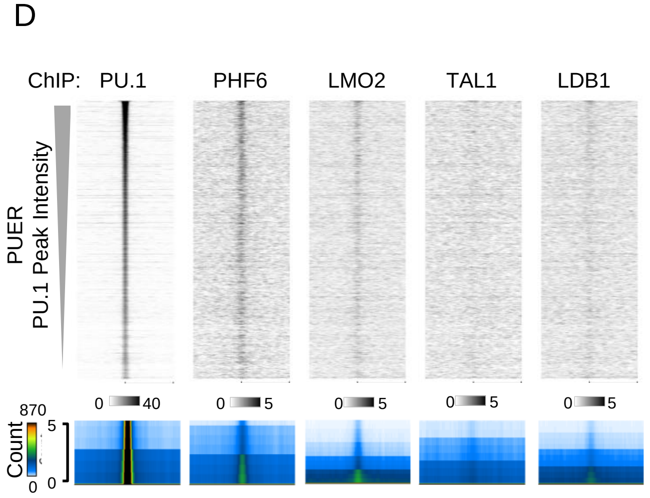


E

**GO for Peak Associations**

| **PHF6-L/T/L peaks** | ***p*-value** |
|---|---|
| positive regulation of GTPase activity | 1.22E-06 |
| regulation of GTPase activity | 2.78E-06 |

| **PHF6-only peaks** | ***p*-value** |
|---|---|
| phagocytosis | 3.20E-06 |
| nuclear-transcribed mRNA catabolic process | 3.84E-06 |
| mRNA catabolic process | 4.06E-06 |
| protein N-linked glycosylation | 5.96E-05 |
| positive regulation of myeloid cell differentiation | 2.33E-04 |

F

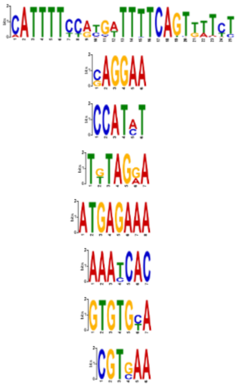

| Motifs Enriched in PHF6-L/T/L peaks | *p*-value | Name | /282 |
|---|---|---|---|
| | 3.4e-471 | NFAT | 64 |
| | 9.1e-019 | ETS | 134 |
| | 9.4e-015 | E-box | 96 |
| | 3.0e-014 | | 63 |
| | 7.9e-012 | IRX3 | 59 |
| | 1.6e-011 | GFI1 | 61 |
| | 8.1e-005 | ZSCAN4 | 44 |
| | 9.9e-005 | PAX6 | 52 |

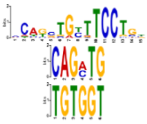

| Motifs Enriched in PHF6-only peaks | *p*-value | Name | /589 |
|---|---|---|---|
| | 1.0e-120 | E-box/ETS1 | 138 |
| | 2.8e-005 | E-box | 164 |
| | 4.6e-003 | RUNX | 98 |

### Figure S4

A

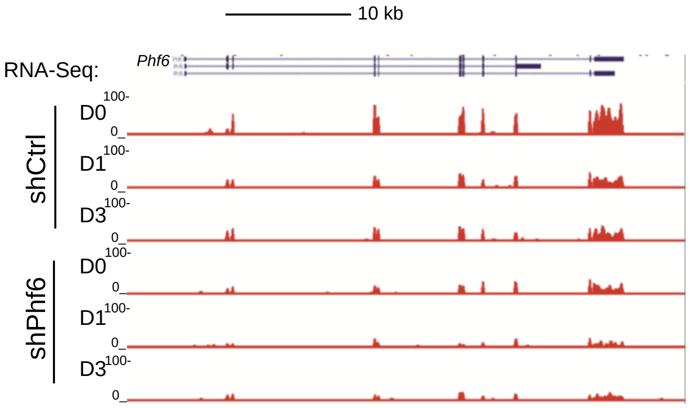


B

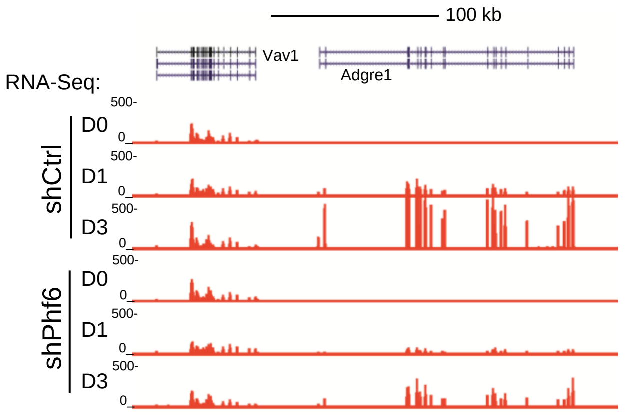


C

| D0 | shCtrl | r_shCtrl | shPhf6 | r_shPhf6 |
|---|---|---|---|---|
| Average FPKM | 33.96 | 24.09 | 34.88 | 23.30 |
| StDev FPKM | 158.89 | 57.05 | 152.86 | 52.26 |
| *p* | 0.0073 | | 0.0014 | |

| D1 | shCtrl | r_shCtrl | shPhf6 | r_shPhf6 |
|---|---|---|---|---|
| Average FPKM | 30.50 | 21.98 | 32.85 | 21.97 |
| StDev FPKM | 184.57 | 58.62 | 210.90 | 58.67 |
| *p* | 0.0328 | | 0.0188 | |

| D3 | shCtrl | r_shCtrl | shPhf6 | r_shPhf6 |
|---|---|---|---|---|
| Average FPKM | 29.85 | 21.68 | 29.71 | 22.78 |
| StDev FPKM | 169.20 | 67.12 | 144.78 | 82.30 |
| *p* | 0.0303 | | 0.04104 | |

Figure S5

A

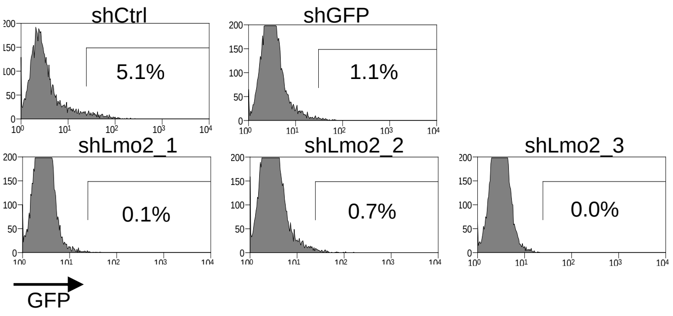
shCtrl
5.1%
shGFP
1.1%
shLmo2_1
0.1%
shLmo2_2
0.7%
shLmo2_3
0.0%
GFP

B

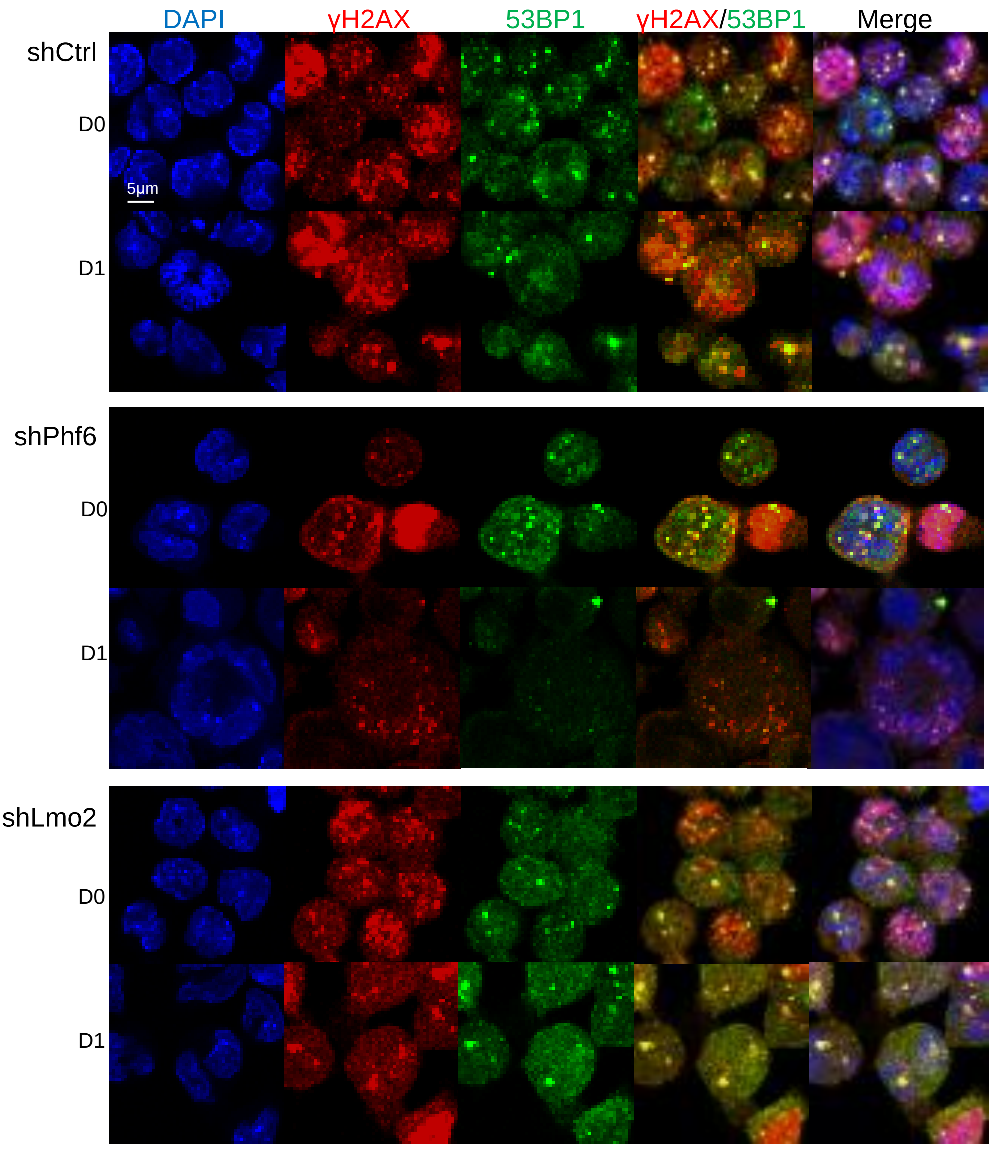
DAPI
γH2AX
53BP1
γH2AX/53BP1
Merge
shCtrl
D0
D1
5μm
shPhf6
D0
D1
shLmo2
D0
D1
